# CTCF knockout in zebrafish induces alterations in regulatory landscapes and developmental gene expression

**DOI:** 10.1101/2020.09.08.282707

**Authors:** Martin Franke, Elisa De la Calle-Mustienes, Ana Neto, Rafael D. Acemel, Juan J. Tena, José M. Santos-Pereira, José L. Gómez-Skarmeta

**Affiliations:** Centro Andaluz de Biología del Desarrollo (CABD), Consejo Superior de Investigaciones Científicas/Universidad Pablo de Olavide, 41013 Seville, Spain

## Abstract

CTCF is an 11-zinc-finger DNA-binding protein that acts as a transcriptional repressor and insulator as well as an architectural protein required for 3D genome folding^1–5^. CTCF mediates long-range chromatin looping and is enriched at the boundaries of topologically associating domains, which are sub-megabase chromatin structures that are believed to facilitate enhancer-promoter interactions within regulatory landscapes ^6–12^. Although CTCF is essential for cycling cells and developing embryos^13,14^, its *in vitro* removal has only modest effects over gene expression^5,15^, challenging the concept that CTCF-mediated chromatin interactions and topologically associated domains are a fundamental requirement for gene regulation^16–18^. Here we link the loss of chromatin structure and gene regulation in an *in vivo* model and during animal development. We generated a *ctcf* knockout mutant in zebrafish that allows us to monitor the effect of CTCF loss of function during embryo patterning and organogenesis. CTCF absence leads to loss of chromatin structure in zebrafish embryos and affects the expression of thousands of genes, including many developmental genes. In addition, chromatin accessibility, both at CTCF binding sites and *cis*-regulatory elements, is severely compromised in *ctcf* mutants. Probing chromatin interactions from developmental genes at high resolution, we further demonstrate that promoters fail to fully establish long-range contacts with their associated regulatory landscapes, leading to altered gene expression patterns and disruption of developmental programs. Our results demonstrate that CTCF and topologically associating domains are essential to regulate gene expression during embryonic development, providing the structural basis for the establishment of developmental gene regulatory landscapes.

---

Vertebrate genomes are folded within the nucleus in a hierarchical manner leading to different levels of chromatin structure that range from chromosome territories to nucleosomes^19–23^. At the Kilo- to Megabases scale, chromatin is organized in topologically associating domains (TADs)^6–9^. According to the current theory, TADs emerge when the cohesin complex, while extruding chromatin, is halted by the CTCF architectural protein^24,25^. Indeed, acute depletion of CTCF or cohesin in cultured cells lead to a severe loss of TAD insulation or the disappearance of all chromatin loops, respectively^5,26^. Recent evidences have suggested that TADs facilitate the contact of *cis*-regulatory elements (CREs) with promoters located within them, while preventing interactions with promoters located in neighboring TADs. In this sense, genomic structural variations that rearrange TAD boundaries lead to enhancer-promoter rewiring, alterations in gene expression and congenital malformations^27–31^. However, to what extent TADs are crucial for gene regulation is currently under debate. Depletion of CTCF in mammalian *in vitro* systems causes only modest transcriptional alterations^5,15,32^, in agreement with some *in vivo* studies^33,34^. However, targeted deletion of several CTCF sites and quantitative gene expression analyses reveal loss of gene expression^35–37^. In fact, our understanding of CTCF function *in vivo* is limited due to its essential function during the cell cycle and the early embryonic lethality in mice^13,14^. Here, we analyze the genome-wide effect of CTCF knockout in developing zebrafish embryos, showing that chromatin structure is essential for the precise regulation of developmental genes and thus provides a scaffold for the establishment of developmental gene regulatory landscapes.

## Generation of a zebrafish *ctcf*^−/−^ zygotic mutant

In order to study the requirement of CTCF in an *in vivo* vertebrate model, we have generated a *ctcf* zygotic knockout mutant in zebrafish. Using CRISPR/Cas9 with two single guide RNAs (sgRNAs), we obtained heterozygous *ctcf*^+/−^ adult individuals carrying a 260-bp deletion, encompassing exons 3 and 4 of the *ctcf* gene that leads to a premature stop codon within exon 4. The expected truncated CTCF protein is depleted of all zinc finger domains, preventing CTCF binding to chromatin. While *Ctcf*^−/−^ zygotic knockout mice die at peri-implantation stages^14^, zebrafish mutants undergo gastrulation and organogenesis and develop normally until pharyngula stages, around 24 hours-post-fertilization (hpf). At 24 hpf, *ctcf*^−/−^ mutants are phenotypically indistinguishable from their heterozygous and wild-type siblings. However, at 48 hpf, *ctcf*^−/−^ embryos showed a clear phenotype that includes pigmentation defects, heart edema and reduced size of head and eyes (Fig. 1a), dying shortly after this stage. Immunofluorescence analysis of wild-type and *ctcf*^−/−^ mutant embryos showed that CTCF protein was absent both at 24 and at 48 hpf (Fig. 1b), while maternal *ctcf* mRNA is detected at least until 75% of epiboly (8 hpf, gastrulation)^38^. This suggests that the late lethality of zebrafish *ctcf*^−/−^ mutants compared to mice might be due to the presence of maternal CTCF protein for a longer time during early embryonic development. Therefore, our *ctcf*^−/−^ zebrafish mutant provides a unique tool to examine the contribution of this protein in genome architecture, gene expression and body plan formation in a vertebrate model system. We therefore exploited this model using a combination of chromosome conformation capture, transcriptomic and epigenomic techniques.

**Figure 1.**
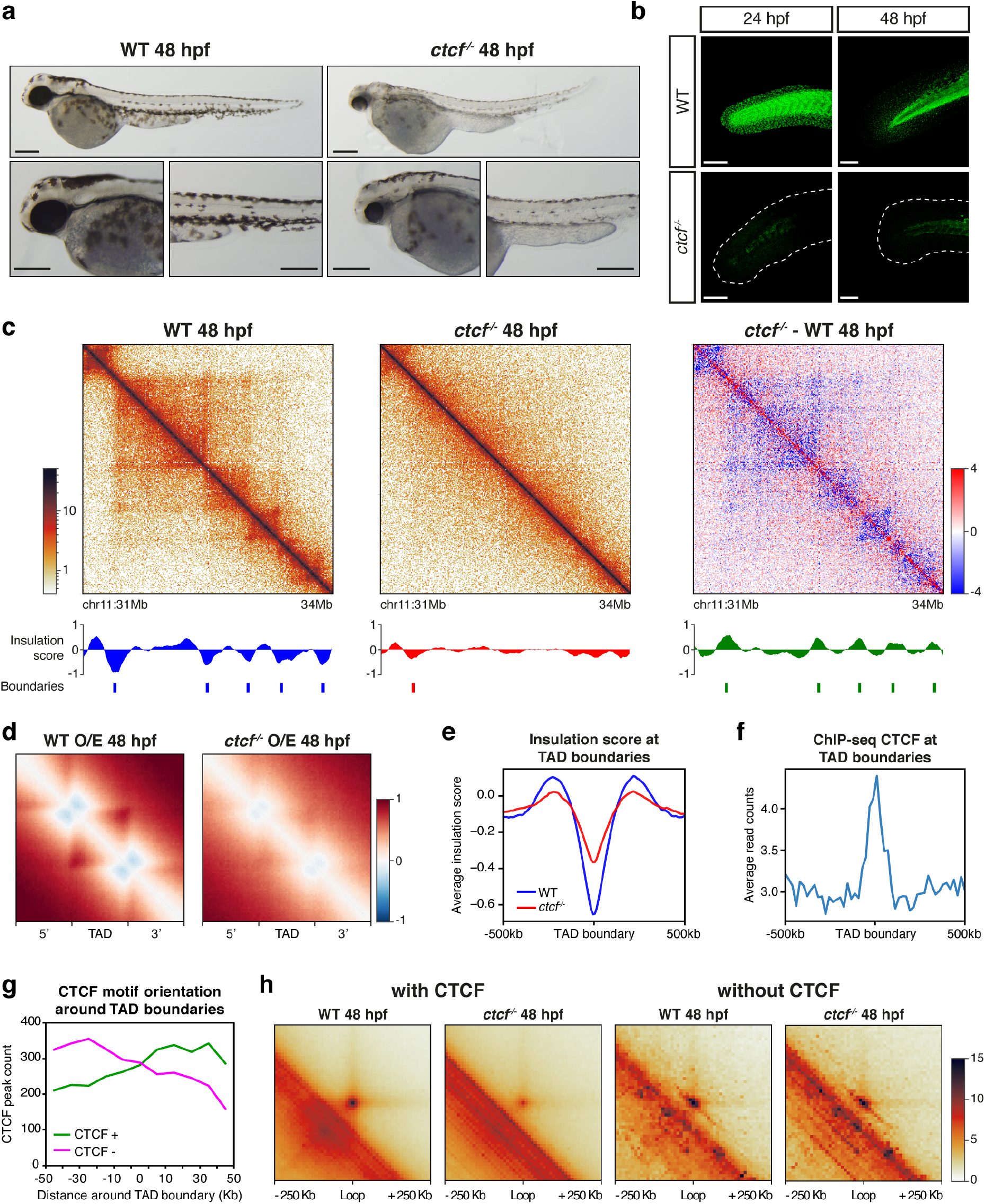
Knockout of *ctcf* in zebrafish embryos disrupts chromatin structure. **a,** Pictures of wild-type (WT) and *ctcf*^−/−^ zebrafish embryos at 48 hours post fertilization (hpf) showing mutant phenotypes, including reduced size of head and eyes, heart edema and defective pigmentation. Scale bars represent 250 μm. **b,** Whole-mount embryo immunofluorescence of CTCF in WT and *ctcf*^−/−^ zebrafish embryos at 24 and 48 hpf showing the absence of this protein in the tail and fin fold in knockout mutants. Scale bars represent 100 μm. **c,** HiC normalized contact maps at 10 Kb resolution from WT and *ctcf*^−/−^ zebrafish embryos, as well as the difference between them, at 48 hpf. A 3-Mb genomic region is plotted, aligned with the insulation scores and the called topologically associating domain (TAD) boundaries. **d,** Aggregate analysis of observed/expected HiC signal in WT and *ctcf*^−/−^ embryos at 48 hpf for the 2,438 TADs called in WT embryos, rescaled and surrounded by windows of the same size. **e,** Average insulation score profiles of WT and *ctcf*^−/−^ zebrafish embryos at 48 hpf around the TAD borders called in the WT. **f,** Average CTCF ChIP-seq signal in WT embryos at 48 hpf around TAD boundaries. **g,** CTCF peak count of those peaks containing CTCF motifs located in the positive (CTCF +) or negative (CTCF −) strands around TAD boundaries, showing a clear preference for CTCF + motifs in the 3’ side of the boundary and for CTCF − motifs in the 5’ side of the boundary. **h,** Aggregate peak analysis of chromatin loops called by HiCCUPs with or without CTCF binding at 48 hpf.

## CTCF is required for chromatin organization in zebrafish embryos

We first analyzed whether the absence of CTCF in zebrafish embryos caused loss of chromatin structure, as previously reported in *in vitro* models^5,15^. For this, we performed HiC experiments in wild-type and *ctcf*^−/−^ whole embryos at 48 hpf and visualized the data at 10-Kb resolution. Figure 1c shows that chromatin structure was established at this stage in wild-type embryos, similar to previous reports^39^, detecting 2,438 TADs based on insulation scores^40^. Other 3D chromatin features commonly detected at this scale, such as loops and stripes, were also observed. In contrast, we found a general loss of chromatin structure in *ctcf*^−/−^ embryos, leading to the detection of only 1,178 TADs and to a reduction of intra-TAD contacts and insulation in wild-type TADs (Fig. 1c-e; Extended Data Fig. 1a-d). These data confirmed that CTCF is essential for 3D chromosome organization in zebrafish embryos, as described for other vertebrates including mammals and frogs^5,15,41^. Next, we analyzed A and B compartments in wild-type and *ctcf*^−/−^ embryos and, although we found a similar distribution of AB compartments, we detected increased AB interactions and decreased compartmentalization strength in the mutants (Extended Data Fig. 1e-g). This contrasts with previous data in cultured cells^5^ and suggests that CTCF may be required for higher order chromatin structure at least in this *in vivo* context.

We then profiled CTCF binding to chromatin in zebrafish embryos using ChIPmentation and found that wild-type TAD boundaries were enriched for CTCF binding, 97% of them containing CTCF sites (Fig. 1f; Extended Data Fig. 2a-b). In addition, the consensus motifs of CTCF at these binding sites around TAD boundaries were preferentially located in a convergent orientation (Fig. 1g), consistent with previous observations^6,10,39,42,43^. Next, we called chromatin loops in wild-type embryos and detected 1,297 loops, 90% of which contained CTCF binding sites at least at one of the anchors (Extended Data Fig. 2). Interestingly, aggregate peak analysis of CTCF-containing chromatin loops showed a marked decrease in intensity in *ctcf*^−/−^ mutants, while 10% of loops without CTCF remained largely unaffected by CTCF loss (Fig. 1h), suggesting that they may be formed by CTCF-independent mechanisms. Therefore, we conclude that CTCF is essential for the establishment of most chromatin loops in zebrafish embryos, similarly to other vertebrates^5,41^.

## Developmental gene expression requires CTCF

To analyze the effects of CTCF absence over gene expression *in vivo*, we performed RNA-seq on whole embryos at 24 and 48 hpf. At 24 hpf, we detected 260 up- and 458 down-regulated genes (Fig. 2a). However, at 48 hpf, we detected as much as 2,730 up- and 3,324 down-regulated genes (Fig. 2b). Strikingly, while differentially expressed genes (DEGs) at 24 hpf were enriched only in biological functions related to immune and DNA damage responses, DEGs at 48 hpf were enriched, among other general functions, in transcription regulation and developmental processes including skeletal muscle development or nervous system development (Fig. 2c-d). This indicates that CTCF is required for the expression of thousands of genes during zebrafish development, an impacting result that contrasts to previous observations in *in vitro* experimental setups showing alteration of a few hundred genes upon CTCF removal^5,32^. Next, we analyzed gene expression changes in the transition from 24 to 48 hpf in wild-type embryos and found that genes that get activated in this period tend to be down-regulated in *ctcf*^−/−^ embryos, and vice versa, indicating that many developmental genes fail to acquire their normal expression level during this developmental period (Extended Data Fig. 3).

**Figure 2.**
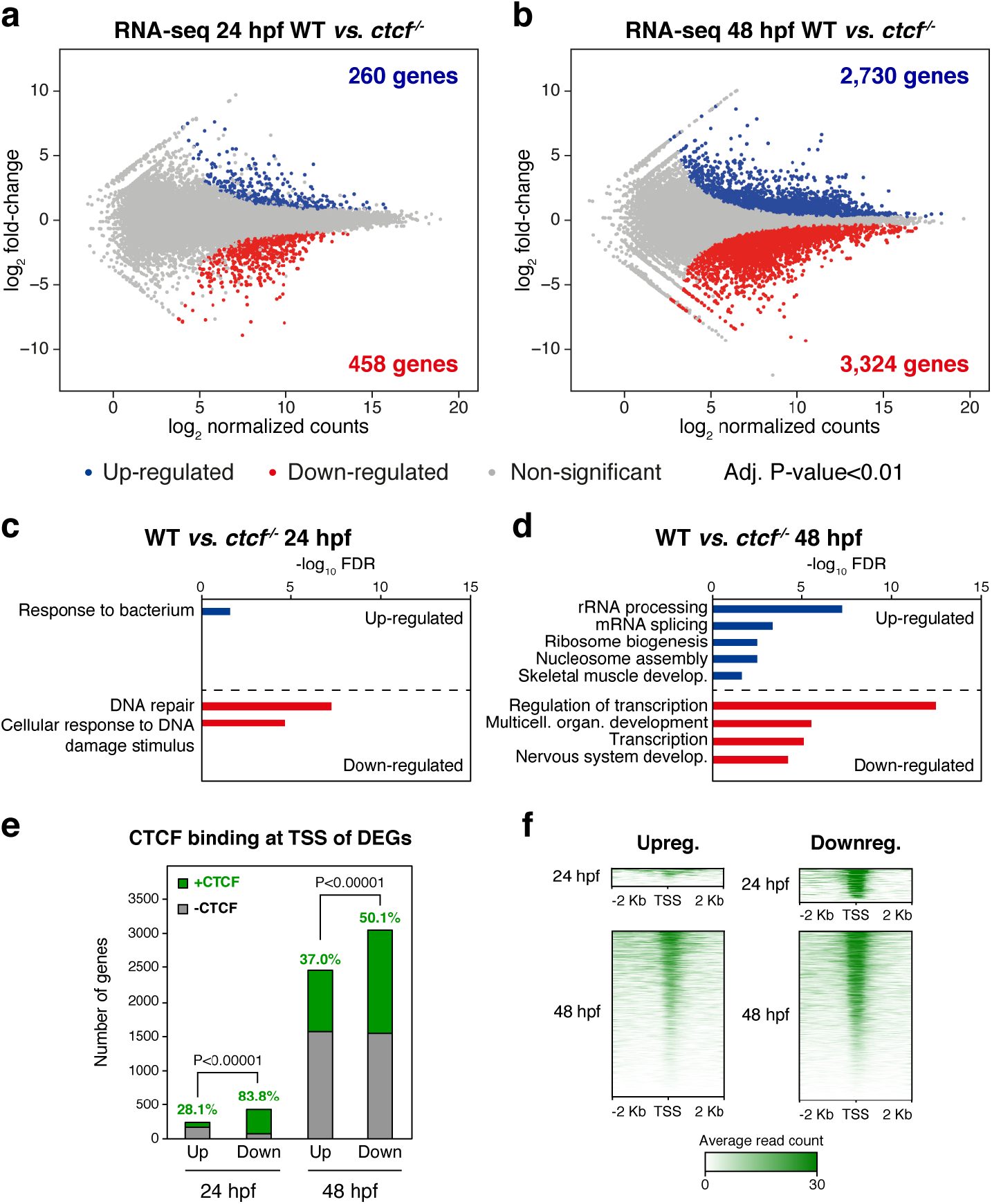
CTCF absence in zebrafish embryos leads to altered developmental gene expression. **a-b,** Differential analyses of gene expression between WT and *ctcf*^−/−^ embryos at 24 (a) and 48 hpf (b) from RNA-seq data (n = 2 biological replicates per condition). The log_2_ normalized read counts of WT transcripts versus the log_2_ fold-change of expression are plotted. Transcripts showing a statistically significant differential expression (adjusted P-value < 0.01) are highlighted in blue (up-regulated) or red (down-regulated). The number of genes that correspond to the up- and down-regulated transcripts are shown inside the boxes. **c-d,** Gene Ontology (GO) enrichment analyses of biological processes for up- and down-regulated genes in *ctcf*^−/−^ embryos at 24 (c) and 48 hpf (d). Terms with a false discovery rate (FDR) < 0.05 are shown and considered as enriched. **e,** Number of differentially expressed genes (DEGs) at 24 and 48 hpf showing (green) or not (grey) CTCF binding at their transcription start sites (TSSs). **f,** Heatmaps showing CTCF ChIP-seq signal around the TSS of DEGs at 24 and 48 hpf.

We then explored the possible function of CTCF to directly regulate DEGs by analyzing its binding to their transcription start sites (TSSs). At 24 hpf, we found a clear bias of CTCF binding towards the TSS of down-regulated genes (83.8%) as compared to up-regulated genes (28.1%) (Fig. 2e-f). This confirms previous observations^5,15^ and suggests distinct mechanisms of CTCF function at activated and repressed genes. By contrast, only 50.1% of down-regulated and 37.0% of up-regulated genes at 48 hpf showed CTCF binding at their TSSs (Fig. 2e-f). Interestingly, we observed that down-regulated genes that are enriched in developmental functions were mainly those without CTCF bound at their TSSs (Extended Data Fig. 4), raising the possibility that developmental genes could be de-regulated indirectly due to defects in chromatin folding. Altogether, these data show that CTCF absence leads to altered developmental gene expression that may account for the observed developmental abnormalities.

## CTCF is required for chromatin accessibility at developmental CREs

The expression of developmental genes is often regulated by multiple tissue-specific CREs, on which combinations of transcription factors (TFs) are bound, giving rise to precise spatial and temporal expression patterns. Since CTCF absence affects the expression of developmental genes mostly without binding to their promoters, we reasoned that this could be due to alterations in the function of their associated CREs. To test this, we performed ATAC-seq in wild-type and *ctcf*^−/−^ embryos at 24 and 48 hpf. At 24 hpf, we only found 56 differentially accessible regions (DARs), 21 with increased (up-regulated) and 35 with decreased accessibility (down-regulated) (Fig. 3a). However, at 48 hpf we found a total of 18,744 DARs, most of them down-regulated (18,138 sites vs. 606 up-regulated) (Fig. 3b), temporally coinciding with the detected altered expression of developmental genes (Fig. 2b). Indeed, when we analyzed CREs gaining or losing accessibility in wild-type embryos from 24 to 48 hpf, we found that these sites failed to gain or lose accessibility in *ctcf*^−/−^ embryos (Extended Data Fig. 5), indicating that loss of CTCF impacts chromatin accessibility of thousands of CREs. Motif enrichment analysis showed that the CTCF consensus binding sequence was specifically enriched in down-regulated peaks, both at 24 and 48 hpf (Fig. 3c-d). We confirmed this by analyzing CTCF binding to DARs at 48 hpf and found that 17.5% of up-regulated but 53.5% of down-regulated peaks were bound by CTCF (Fig. 3e). In contrast, up-regulated peaks were enriched for the p53 family motif at 48 hpf. At this stage we also found increased expression of *tp53* and well-known p53 target genes (Fig. 3d; Extended Data Fig. 6), pointing towards an increased apoptotic response in *ctcf*^−/−^ mutants^14^. To test a possible contribution of p53 to the mutant phenotypes, we injected one-cell stage embryos with a morpholino to knock-down *tp53* expression. Despite reduced p53-target gene expression and loss of p53-target motif in morpholino-injected mutants, differential accessibility remained unaffected (Extended Data Fig. 6). Furthermore, the p53 knockdown did not change the mutant phenotype at 48 hpf, indicating that the phenotypic response is not driven by pro-apoptotic processes.

Next, we associated DARs to nearby DEGs and found that the average change in gene expression was consistent with the tendency of changes in chromatin accessibility and independent of CTCF binding (Fig. 3f). Interestingly, only down-regulated peaks without CTCF binding were associated with genes enriched in developmental functions, such as hindbrain development or heart formation (Fig. 3g). This indicates that loss of CTCF affects indirectly the accessibility of developmental CREs. We also noted that down-regulated ATAC peaks without CTCF binding sites were highly clustered within the regulatory landscapes of developmental genes, many of them strongly down-regulated in the mutant (Extended Data Fig. 7a-c). This is consistent with the view that developmental genes frequently locate within large gene deserts containing many CREs. Indeed, we found that TADs containing miss-regulated developmental genes were larger and had more associated CREs than those containing non-developmental genes (Fig. 3h-i). Several examples illustrate this tendency. The *sall1a* gene, encoding a transcriptional repressor involved in organogenesis, is in a TAD whose structure was lost in *ctcf*^−/−^ embryos (Fig. 3j). The expression of *sall1a* was reduced in the absence of CTCF and several CREs exhibited reduced accessibility with most of them not binding CTCF. Other examples included the *lhx1a* and *sox11b* genes, both encoding developmental transcription factors (Extended Data Fig. 7d-e). Altogether, these data show that CTCF is required for the accessibility of thousands of CREs, many of which are associated with developmental genes.

**Figure 3.**
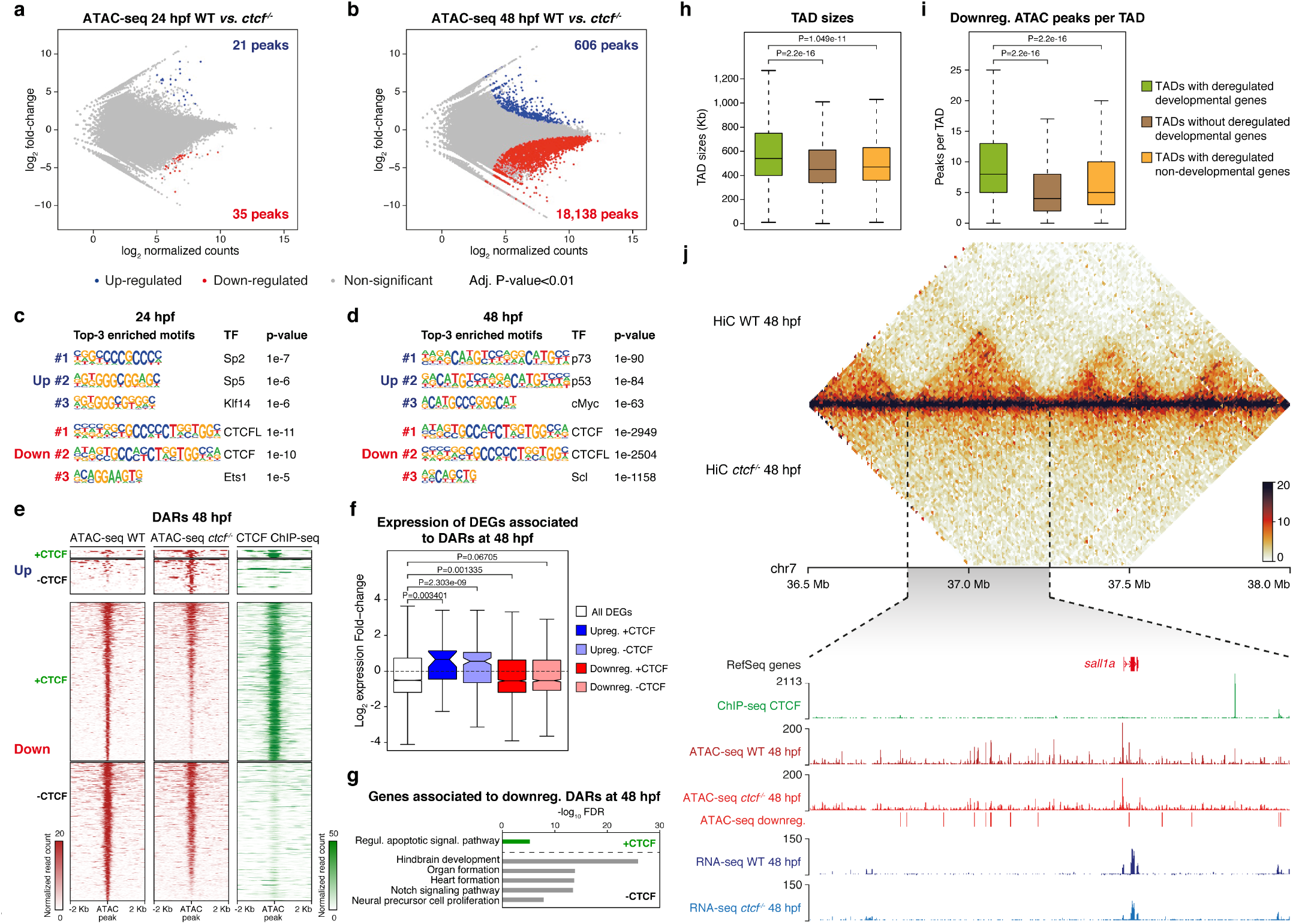
CTCF promotes chromatin accessibility at developmental *cis*-regulatory elements. **a-b,** Differential analyses of chromatin accessibility between WT and *ctcf*^−/−^ embryos at 24 (a) and 48 hpf (b) from ATAC-seq data (n = 2 biological replicates per condition). The log_2_ normalized read counts of WT ATAC peaks versus the log_2_ fold-change of accessibility are plotted. Regions showing a statistically significant differential accessibility (adjusted P-value < 0.01) are highlighted in blue (up-regulated) or red (down-regulated). The number of peaks that correspond to the up- and down-regulated sites are shown inside the boxes. **c-d,** Motif enrichment analyses for the up- and down-regulated ATAC peaks in *ctcf*^−/−^ embryos at 24 (c) and 48 hpf (d). The 3 motifs with the lowest p-values are shown for each case. **e,** Heatmaps plotting normalized ATAC-seq signal in WT and *ctcf*^−/−^ embryos at 48 hpf (red), as well as CTCF ChIP-seq signal (green), for the differentially accessible regions (DARs) from (b) overlapping or not with CTCF peaks. **f,** Box plots showing the expression fold-change in *ctcf*^−/−^ embryos at 48 hpf of all DEGs or only those associated with up-regulated or down-regulated DARs, overlapping or not with CTCF sites. Center line, median; box limits, upper and lower quartiles; whiskers, 1.5x interquartile range; notches, 95% confidence interval of the median. Statistical significance was assessed using the Wilcoxon’s rank sum test. **g,** GO enrichment analyses of biological processes for the genes associated with down-regulated DARs in *ctcf*^−/−^ embryos at 48 hpf, overlapping or not with CTCF sites. GO terms showing an FDR < 0.05 are considered as enriched. **h-i,** Box plots showing the TAD sizes (h) and the number of down-regulated DARs per TAD (i) for TADs containing developmental miss-regulated genes, TADs not containing developmental miss-regulated genes and TADs containing only non-developmental miss-regulated genes. Center line, median; box limits, upper and lower quartiles; whiskers, 1.5x interquartile range. Statistical significance was assessed using the Wilcoxon’s rank sum test. **j,** Top, heatmaps showing HiC signal in WT and *ctcf*^−/−^ embryos at 48 hpf in a 1.5-Mb region of chromosome 7. Bottom, zoom within the *sall1a* TAD showing UCSC Genome Browser tracks with CTCF ChIP-seq, ATAC-seq at 48 hpf in WT and *ctcf*^−/−^ embryos, ATAC-seq down-regulated peaks and RNA-seq at 48 hpf in WT and *ctcf*^−/−^ embryos. The *sall1a* gene is shown in red because it is down-regulated.

## CTCF is required for the spatiotemporal expression patterns of developmental genes

We have shown so far that CTCF is not only required for chromosome folding during zebrafish development, but also for the robust expression levels of many developmental genes and chromatin accessibility at their regulatory landscapes. To better assess gene miss-expression in relation to the loss of chromatin structure, we first investigated chromatin interactions at the enhancer-promoter level with high resolution by performing UMI-4C experiments. We used developmental gene promoters as viewpoints to analyze their regulatory landscapes in wild-type and *ctcf*^−/−^ embryos at 48 hpf, such as the *ptch2* promoter. *Ptch2* is a patterning gene that encodes a cell receptor binding the Shh morphogen and whose expression was detected as up-regulated by RNA-seq in *ctcf*^−/−^ mutants (Fig. 4a). We found that contacts from the *ptch2* promoter spanned a region of about 500 kb in wild-type embryos, establishing contacts with many genomic regions that included ATAC peaks (potential CREs) with and without CTCF binding (Fig. 4a). However, the *ptch2* regulatory landscape was drastically reduced in *ctcf*^−/−^ embryos. The interaction profile was generally characterized by a loss of long-range contacts but retaining some contacts at shorter ranges (Fig. 4a), consistent with observations in mammalian cells^32,44^. Genomic regions showing a reduced contact frequency with the *ptch2* promoter included CTCF-binding sites as well as ATAC peaks with reduced accessibility in the mutant and others not affected by CTCF absence. These results indicate that enhancer-promoter contacts were severely affected by the absence of CTCF, and in particular, long-range interactions.

**Figure 4.**
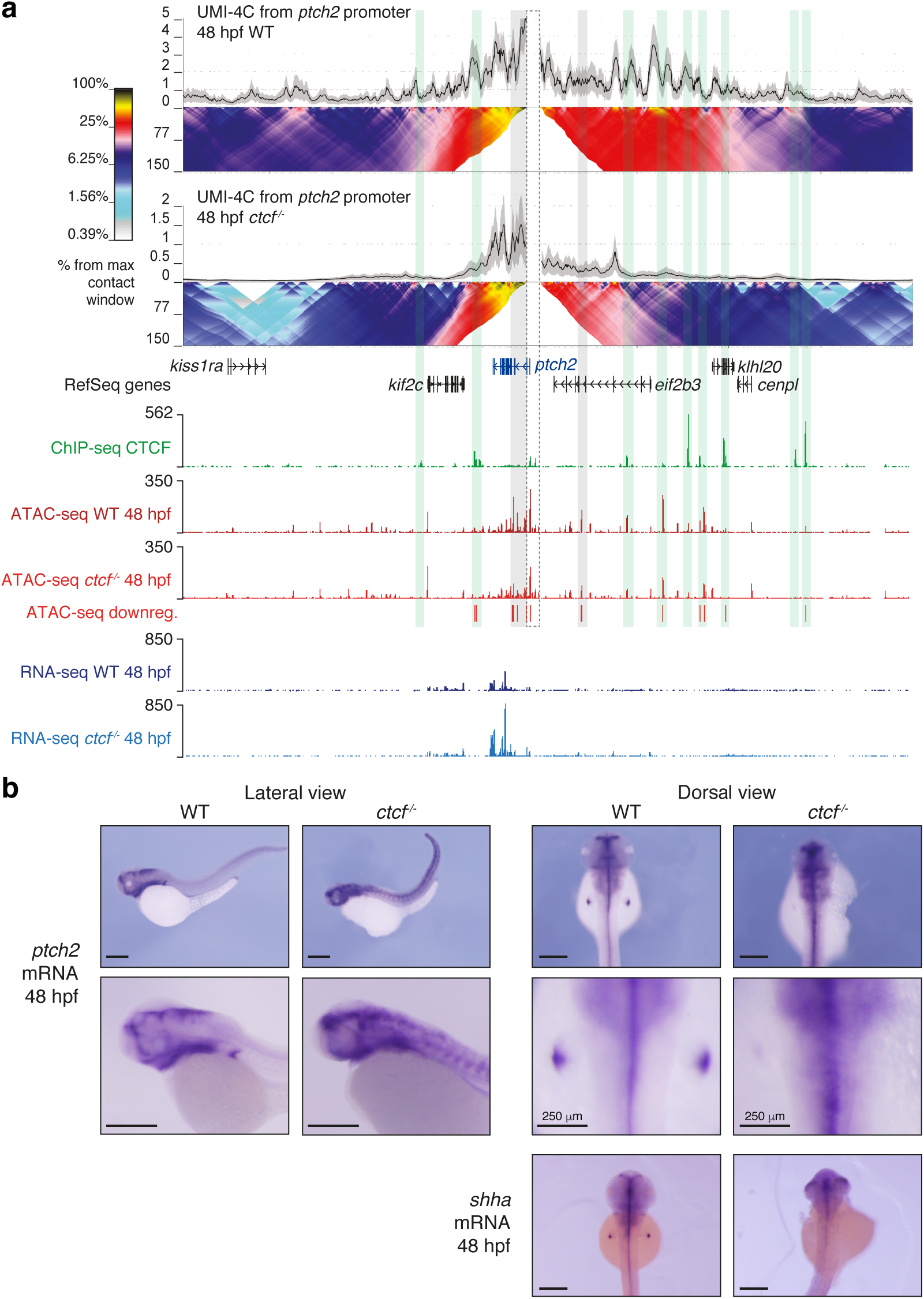
CTCF is required to sustain the regulatory landscapes and complex expression patterns of developmental genes. **a,** Top, UMI-4C assays in WT and *ctcf*^−/−^ embryos at 48 hpf using the *ptch2* gene promoter as a viewpoint. Black lines and grey shadows represent the average normalized UMI counts and their standard deviation, respectively. Domainograms below UMI counts represent contact frequency between pairs of genomic regions. Bottom, UCSC Genome Browser tracks with CTCF ChIP-seq, ATAC-seq at 48 hpf in WT and *ctcf*^−/−^ embryos, ATAC-seq down-regulated peaks and RNA-seq at 48 hpf in WT and *ctcf*^−/−^ embryos. The *ptch2* gene is shown in blue because it is up-regulated. A dotted-line square represents the restriction fragment containing the *ptch2* gene promoter that is used as a viewpoint; green shadows highlight CTCF sites and grey shadows highlight down-regulated ATAC-peaks without CTCF binding. **b,** Whole-mount *in situ* hybridization of the *ptch2* and *shha* genes in WT and *ctcf*^−/−^ embryos at 48 hpf. Left, lateral view; right, dorsal view. Scale bars represent 500 μm, unless indicated.

Next, we investigated whether this loss of contacts altered the expression pattern of *ptch2* by performing whole-embryo *in situ* hybridization. *Ptch2* mRNA was detected in the brain, pharyngeal arches and pectoral fin buds of wild-type embryos, but we found that this pattern was severely altered in *ctcf*^−/−^ embryos (Fig. 4b). Consistent with the upregulation in our bulk RNA-seq data, *ptch2* expression in mutant embryos was extended to broader regions of the brain, pharyngeal arches, neural tube and a prominent expansion of expression was observed in the somites; however, expression in the pectoral fin buds was lost (Fig. 4b), illustrating the limitation of bulk RNA-seq to detect complex changes in gene expression patterns. Consistently, a similar effect was found for expression of *shha* in the pectoral fin buds (Fig. 4b). These results indicate that loss of CTCF and promoter contacts alters the expression patterns of developmental genes, including loss and gain of expression domains, likely disrupting developmental programs. Similar changes in the chromatin interactions of regulatory landscapes and gene expression patterns were observed at the HoxD cluster. Viewpoints from the promoters of *hoxd4a* and *hoxd13a* showed reduced interactions within their regulatory landscapes in *ctcf*^−/−^ embryos, especially long-range contacts (Extended Data Fig. 8a). Although we could not detect mis-regulation of *hoxd4a* and *hoxd13a* by RNA-seq, *in situ* hybridization experiments showed a clear reduction of their expression levels (Extended Data Fig. 8b). However, other *hox* genes showed consistent mis-regulation detected by both techniques (Extended Data Fig. 8c). Altogether, our data indicate that CTCF is required to establish chromatin contacts of gene promoters with their associated regulatory landscape and to ensure the accurate spatiotemporal expression patterns of developmental genes.

## Discussion

In this work, we have established a new *in vivo* model to study the loss of CTCF in zebrafish and demonstrate that chromatin structure is required to maintain developmental gene regulatory landscapes during body plan formation. In the last years, the function of CTCF in chromosome folding has been clearly demonstrated in mammalian *in vitro* systems, including mouse embryonic stem cells, neural progenitor cells as well as human morula embryos^5,15,32^. These studies showed by different depletion mechanisms that CTCF knock-down severely reduces TAD formation and insulation. Accordingly, we show here that CTCF is also required for chromatin structure in zebrafish embryos (Fig. 1), extending these conclusions to vertebrates and in agreement with a recent report showing that CTCF knockdown in *Xenopus* embryos altered chromatin structure^41^.

Despite this well-known function of CTCF, its requirement for the regulation of gene expression has remained controversial. The studies mentioned above showed modest effects of CTCF depletion in gene expression, suggesting that steady-state transcription is mostly resistant to genome-wide alteration of chromatin structure. This contrasts with the observation that CTCF is essential for embryonic development^14^, but suggests that CTCF-mediated chromatin structure could be essential for processes in which cells respond to multiple signals and where transcriptional control is highly dynamic. However, the early embryonic lethality of CTCF knockout in animal models, has impeded the analysis of CTCF function for transcriptional regulation beyond pluripotency. Our *ctcf* mutant zebrafish model overcame this limitation due to the prolonged maternal contribution that lasts, at least, until gastrulation. This allows *ctcf*^−/−^ embryos to develop until stages in which patterning and organogenesis take place. Using this model, we observe for the first time in developing embryos the miss-regulation of thousands of genes (Fig. 2), among which many lineage-specific genes that are dynamically regulated during development. These observations are consistent with recent reports, showing that CTCF is required for the expression of a subset of lineage-specific genes during cell differentiation^32^ and for fast transcriptional responses to external stimuli^45^.

The expression of developmental genes is characterized by a tight spatiotemporal control by CREs that constitute their regulatory landscapes. These have been shown to largely coincide with TADs and to be constrained by TAD boundaries^46^. Here, we show that chromatin accessibility at CTCF sites but also at thousands of CREs is compromised in *ctcf* mutants (Fig. 3). Specifically, clusters of CREs within large TADs of developmental genes show highly reduced accessibility, most of them without direct CTCF binding, suggesting an indirect effect due to the loss of chromatin structure. This may arise because of reduced TF accumulation at CREs either due to their decreased expression levels or decreased enhancer-promoter interactions. High-resolution analyses of the *ptch2*, *hoxd4a* and *hoxd13a* gene regulatory landscapes show that CREs with reduced accessibility lose contacts with their promoters, mainly long-range and many of them without CTCF binding (Fig. 4 and Extended Data Fig. 8). While it is unlikely that CTCF directly mediates those enhancer-promoter interactions, it may favor their establishment by promoting contacts within the involved TADs. This is in agreement with recent observations showing that CTCF is required for long-range enhancer-promoter contacts^32,44^. Consequently, the complex expression patterns of these genes are altered in a tissue-specific manner, showing up- and down-regulation in different embryonic domains (Fig. 4). This whole-embryo *in situ* hybridization approach allows the detection of transcriptional alterations not detected by bulk RNA-seq, highlighting the potential of using animal models *versus in vitro* systems.

In summary, our data demonstrate that CTCF is essential to sustain large regulatory landscapes of developmental genes during embryonic development. This would favor the proper interaction of multiple CREs with their target genes, leading to the complex spatiotemporal expression patterns of developmental genes. It has been suggested that TADs may have evolved as conserved scaffolds for developmental gene regulatory landscapes^47^. Our observations support this view by linking chromatin structure at regulatory landscapes with gene function.

## Methods

### Animal experimentation

Wild-type AB/Tübingen zebrafish strains were maintained and bred under standard conditions. All experiments involving animals conform national and European Community standards for the use of animals in experimentation and were approved by the Ethical Committees from the University Pablo de Olavide, CSIC and the Andalusian government.

### CRISPR-Cas9 genome editing

CRISPR target sites to mutate the *ctcf* gene were identified using the CRISPRscan online tool^48^. Two single guide RNAs (sgRNAs) targeting the exons 4 and 5 of the *ctcf* gene were used with the following target sequences: 5’-GGA GTT ACA CTT GCC CAC GC-3’ and 5’-GGC ATG GCC TTT GTC ACC AG-3’. The template DNA for sgRNA transcription was generated by PCR using CTCFexon4, CTCFexon5 and sgRNA_universal primers (Extended Data Table 1) and Phusion DNA polymerase (Thermo Fisher Scientific). sgRNAs were *in vitro* transcribed using the HiScribe T7 Quick High Yield RNA synthesis kit (NEB) using 75 ng of template, treated with DNase I (NEB) and purified using the RNA Clean and Concentrator kit (Zymo Research).

One-cell stage zebrafish embryos were injected with 2-3 nl of a solution containing 140 ng/*μ*l of Cas9 mRNA and 25 ng/*μ*l of each sgRNA. The CRISPR-Cas9 approach generated a deletion of 260 bp encompassing exons 4 and 5 and resulting in a premature STOP codon in exon 5. The predicted truncated protein had 343 amino acids instead of 798, lacking ten and a half of the eleven zinc finger domains of the CTCF protein. For genotyping, genomic DNA was obtained by incubating the samples (whole embryos or adult caudal fin fragments) in TE buffer supplemented with 5% Chelex-100 (BioRad) and 10 *μ*g/ml Proteinase K (Roche) for 1h (embryos) or 4h (fins) at 55°C and 10 min at 95°C, and then stored at 4°C. One microliter of the supernatant was used as a template for standard 20 *μ*l PCR reactions using CTCFpF and CTCFpR primers (Extended Data Table 1), resulting in 842- or 582-bp amplicons for wild type or mutant alleles, respectively. The mutant allele was stably maintained in heterozygosis with no apparent phenotypes but homozygous mutants are embryonic lethal (<3 days).

### Whole-mount embryo immunofluorescence

For immunofluorescence, embryos were fixed overnight at 4°C with 4% paraformaldehyde, washed in PBT (PBS supplemented with 0.2% Triton-X100) and blocked in this solution with 2% goat serum and 2 mg/ml BSA for 1 h at RT. Then, they were incubated overnight at 4°C with primary antibody specific for zebrafish CTCF^49^ (used in 1:500 dilution). After extensive washings with PBT, embryos were incubated overnight at 4°C with goat anti-rabbit Alexa Fluor 488 secondary antibody (used 1:800 dilution, A27034 Invitrogen). Finally, embryos were flat-mounted and imaged under an SP confocal microscope (Leica).

### Whole-mount embryo *in situ* hybridization

Antisense RNA probes were prepared from cDNA using digoxigenin (Boehringer Mannheim) as label and the primers listed in Extended Data Table 1, except those for *shha* and *hoxd13a* that were previously described^50^. Zebrafish embryos were prepared, hybridized and stained using standard protocols^51^. Embryos at 48 hpf stage were fixed in 4% paraformaldehyde overnight, dehydrated in methanol and stored at −20°C. All solutions and reagents used were RNAse-free. The embryos were hydrated using decreasing amounts of methanol and finally in PBS-0.1% Tween. Then, they were treated with 10 *μ*g/ml proteinase K for 10 min at room temperature and gently washed with PBS-0.1% Tween. In the pre-hybridization step, embryos were kept at 70°C in the hybridization buffer for at least 1 hour. Then, the probe was diluted to 2 ng/*μ*l in hybridization buffer and incubated overnight at 70°C while moving. Pre-heated buffers with decreasing amounts of hybridization buffer (75%, 50%, 25% and 0%) in 2x SSC solution were used to wash embryos for 10 min, plus a 30 min wash at 70°C with 0.05x SSC. Then, they were incubated with Blocking Buffer (PBS-0.1% Tween, 2% normal goat serum, 2 mg/ml bovine serum albumin [BSA]) for 1 hour, and with an anti-digoxigenin antibody (1:5,000 in Blocking Buffer) for at least 2 hours at room temperature. After this, embryos were washed six times with PBS-0.1% Tween at room temperature and then overnight at 4°C. Next day, embryos were washed once more with PBS-0.1% Tween and three times with fresh AP buffer (100 mM Tris-HCl pH 9.5, 50mM MgCl_2_, 100mM NaCl, 0.1% Tween), followed by signal development with NBT/BCIP solution (225 *μ*g/ml NBT, 175 *μ*g/ml BCIP) in multi-well plates in the dark. Signal development was stopped by washing with PBS-0.1% Tween and fixing with 4% paraformaldehyde. Imaging of the *in situ* hybridization signal was performed in MZ-12 dissecting scope (Leica).

### RNA-seq

For total RNA extraction, wild-type and *ctcf*^−/−^ single embryos at 24 or 48 hpf were collected, manually de-chorionated and suspended in TRIsure (Bioline) with chloroform. DNA was used for genotyping and single wild-type and *ctcf*^−/−^ individuals were selected for RNA-seq experiments. Precipitated RNA was then treated with TURBO DNA free kit (Invitrogen). Two biological replicates were used for each analyzed genotype and stage.

Illumina libraries were constructed and sequenced in a BGISEQ-500 single-end lane producing around 50 million (M) of 50-bp reads. Reads were aligned to the GRCz10 (danRer10) zebrafish genome assembly using STAR 2.5.3a^52^ and counted using the htseq-count tool from the HTSeq 0.8.0 toolkit^53^. Differential gene expression analysis was performed using the DESeq2 1.18.1 package in R 3.4.3^54^, setting a corrected P value < 0.01 as the cutoff for statistical significance of the differential expression. Enrichment of GO Biological Process terms was calculated using David^55^, with a false discovery rate (FDR)-corrected P value < 0.05 as statistical cutoff.

### ATAC-seq

ATAC-seq assays were performed using standard protocols^56,57^, with minor modifications. Briefly, single WT or *ctcf*^−/−^ mutant embryos at 24 or 48 hpf coming from *ctcf*^+/−^ crosses were manually de-chorionated. Yolk was dissolved with Ginzburg Ring Finger (55 mM NaCl, 1.8 mM KCl, 1.15 mM NaHCO_3_) by pipetting and shaking 5 min at 1100 rpm. Deyolked embryos were collected by centrifugation for 5 min at 500g 4°C. Supernatant was removed and embryos washed with PBS. Then, embryos were lysed in 50 *μ*l of Lysis Buffer (10 mM Tris-HCl pH 7.4, 10 mM NaCl, 3 mM MgCl_2_, 0.1% NP-40, 1x Roche Complete protease inhibitors cocktail) by pipetting up and down. The whole cell lysate was used for TAGmentation, which were centrifuged for 10 min at 500g 4°C and resuspended in 50 *μ*l of the Transposition Reaction, containing 1.25 *μ*l of Tn5 enzyme and TAGmentation Buffer (10 mM Tris-HCl pH 8.0, 5 mM MgCl2, 10 % w/v dimethylformamide), and incubated for 30 min at 37°C. Immediately after TAGmentation, DNA was purified using the Minelute PCR Purification Kit (Qiagen) and eluted in 20 *μ*l. Before library amplification, purified DNA was used to genotype 24-hpf embryos (see above) and wild-type or *ctcf*^−/−^ mutants were selected for deep sequencing. Libraries were generated by PCR amplification using NEBNext High-Fidelity 2X PCR Master Mix (NEB). The resulting libraries were multiplexed and sequenced in a HiSeq 4000 pair-end lane producing 100M of 49-bp pair end reads per sample.

### ChIPmentation

ChIP-seq of CTCF was performed by ChIPmentation, which incorporates Tn5-mediated TAGmentation of immunoprecipitated DNA, as previously described^58,59^. Briefly, 100 zebrafish embryos at 24 hpf were dechorionated with 300 *μ*g/ml pronase, fixed for 10 min in 1% paraformaldehyde (in 200 mM phosphate buffer) at room temperature, quenched for 5 min with 0.125 M glycine, washed in PBS and frozen at −80°C. Fixed embryos were homogenized in 2 ml cell lysis buffer (10 mM Tris-HCl pH 7.5, 10 mM NaCl, 0.3% NP-40, 1x Roche Complete protease inhibitors cocktail) with a Dounce Homogenizer on ice and centrifuged 5 min 2,300g at 4°C. Pelleted nuclei were resuspended in 333 *μ*l of nuclear lysis buffer (50 mM Tris-HCl pH 7.5, 10 mM EDTA, 1% SDS, 1x Roche Complete protease inhibitors cocktail), kept 5 min on ice and diluted with 667 *μ*l of ChIP dilution buffer (16.7 mM Tris-HCl pH 7.5, 1.2 mM EDTA, 167 mM NaCl, 0.01% SDS, 1.1% Triton-X100). Then, chromatin was sonicated in a Covaris M220 sonicator (duty cycle 10%, PIP 75W, 100 cycles/burst, 10 min) and centrifuged 5 min 18,000g at 4°C. The recovered supernatant, which contained soluble chromatin, was used for ChIP or frozen at −80°C after checking the size of the sonicated chromatin. Four 250 *μ*l aliquots of sonicated chromatin were used for each independent ChIP experiment, and each aliquot incubated with 2 *μ*g of anti-CTCF antibody^49^ and rotated overnight at 4°C. Next day, 20 *μ*l of protein G Dynabeads (Invitrogen) per aliquot were washed twice with ChIP dilution buffer and resuspended in 50 *μ*l/aliquot of the same solution. Immunoprecipitated chromatin was then incubated with washed beads for 1 hour rotating at 4°C and washed twice sequentially with wash buffer 1 (20 mM Tris-HCl pH 7.5, 2 mM EDTA, 150 mM NaCl, 1% SDS, 1% Triton-X100), wash buffer 2 (20 mM Tris-HCl pH 7.5, 2 mM EDTA, 500 mM NaCl, 0.1% SDS, 1% Triton-X100), wash buffer 3 (10 mM Tris-HCl pH 7.5, 1 mM EDTA, 250 mM LiCl, 1% NP-40, 1% Na-deoxycholate) and 10 mM Tris-HCl pH 8.0, using a cold magnet (Invitrogen). Then, beads were resuspended in 25 *μ*l of TAGmentation reaction mix (10 mM Tris-HCl pH 8.0, 5 mM MgCl_2_, 10% w/v dimethylformamide), added 1 *μ*l of Tn5 enzyme and incubated 1 min at 37°C. TAGmentation reaction was put in the cold magnet and the supernatant discarded. Beads were washed twice again with wash buffer 1 and 1x TE and eluted twice for 15 min in 100 *μ*l of elution buffer (50 mM NaHCO3 pH 8.8, 1% SDS). The 200 *μ*l of eluted chromatin per aliquot were then decrosslinked by adding 10 *μ*l of 4M NaCl and 1 *μ*l of 10 mg/ml proteinase K and incubating at 65°C for 6 hours. DNA was purified using Minelute PCR Purification Kit (Qiagen), pooling all aliquots in a single column, and eluted in 20 *μ*l. Library preparation was performed as previously described for ATAC-seq (see above). Libraries were multiplexed and sequenced in a HiSeq 4000 pair-end lane producing around 20M of 49-bp paired-end reads per sample.

### ChIPmentation and ATAC-seq data analyses

ChIPmentation and ATAC-seq reads were aligned to the GRCz10 (danRer10) zebrafish genome assembly using Bowtie2^60^ and those pairs separated by more than 2 Kb were removed. For ATAC-seq, the Tn5 cutting site was determined as the position −4 (minus strand) or +5 (plus strand) from each read start, and this position was extended 5 bp in both directions. Conversion of SAM alignment files to BAM was performed using Samtools^61^. Conversion of BAM to BED files, and peak analyses, such as overlaps or merges, were carried out using the Bedtools suite^62^. Conversion of BED to BigWig files was performed using the genomecov tool from Bedtools and the wigToBigWig utility from UCSC^63^. For ATAC-seq, peaks were called using MACS2 algorithm^64^ with an FDR < 0.05 for each replicate and merged in a single pool of peaks that was used to calculate differentially accessible sites with DESeq2 1.18.1 package in R 3.4.3^54^, setting a corrected P value < 0.01 as the cutoff for statistical significance of the differential accessibility. For ChIPmentation, peaks with an FDR < 0.001 were called with MACS2. For visualization purposes, reads were extended 100 bp for ATAC-seq and 300 bp for ChIPmentation. For data comparison, all ATAC-seq experiments used were normalized using reads falling into peaks to counteract differences in background levels between experiments and replicates, as previously described^58^.

Heatmaps and average profiles of ChIPmentation and ATAC-seq data were generated using computeMatrix, plotHeatmap and plotProfile tools from the Deeptools 2.0 toolkit^65^. TF motif enrichment and peak annotation to genomic features were calculated using the scripts FindMotifsGenome.pl and AnnotatePeaks.pl from Homer software^66^, with standard parameters. For gene assignment to ChIP and ATAC peaks, coordinates were converted to Zv9 (danRer7) genome using the Liftover tool of the UCSC Genome Browser^63^ and assigned to genes using the GREAT tool^67^, with the basal plus extension association rule with standard parameters (5 Kb upstream, 1 Kb downstream, 1 Mb maximum extension). Peak clustering was calculated using the mergeBed tool from Bedtools^62^, considering as clustered those peaks located less than 30 Kb from each other.

### HiC

HiC library preparation was performed as previously described^10^ with minor modifications. Experiments were performed for at least two biological replicates in wild-type and *ctcf*^−/−^ mutant embryos at 48 hpf, using one to three million cells as input material.

#### Embryo fixation and nuclei extraction

Pools of 50 zebrafish embryos were dechorionated with 300 *μ*g/ml pronase, followed by fixation for 10 min in 1% paraformaldehyde (in 200 mM phosphate buffer) at room temperature. The reaction was quenched by adding glycine to a final concentration of 0.125 M and incubation at room temperature for 5 min. Embryos were washed on ice twice with 1x PBS and either snap frozen in liquid nitrogen or processed for nuclei extraction. For nuclei extraction, fixed embryos were homogenized in 2-5 ml freshly prepared lysis buffer (50 mM Tris pH7.5; 150 mM NaCl; 5 mM EDTA; 0.5 % NP-40; 1.15 % Triton X-100; 1x Roche Complete protease inhibitors) with a Dounce Homogenizer on ice. Nuclei were pelleted by centrifugation for 5 min, 750g at 4°C and washed with 1x PBS. Pelleted nuclei were either snap-frozen in liquid nitrogen or further processed.

#### Chromatin digestion

Nuclei pellets were resuspended in 100 *μ*l 0.5% SDS and incubated for 10 min at 62°C, without shaking. 292 *μ*l water and 50 *μ*l 10% Triton X-100 were added to each sample, mixed, and incubated for 15 min at 37°C to quench remaining SDS. 50 *μ*l of 10x restriction enzyme buffer and a total of 400 units of DpnII (NEB, R0543) were added to the sample, mixed and incubated overnight at 37°C with 900 rpm shaking.

#### Biotin fill-in and proximity ligation

Restriction enzyme was heat inactivated. Nuclei were pelleted at 600 g for 10 min at 4°C and resuspended in 445 *μ*l 1x ice-cold NEB buffer 2. For biotin fill-in reaction, 5 *μ*l of 10x NEB buffer 2, 1.5 *μ*l 10 mM (each) dNTP-dATP-mix, 37.5 *μ*l of 0.4 mM biotin-14-dATP and 10 *μ*l of 5 U/ *μ*l Klenow (NEB, M0210L) were added and mixed by pipetting. Samples were incubated at 25°C for 4 h and 800 rpm shaking. To ligate restriction fragment ends, 500 *μ*l of 2x ligation mix (100*μ*l of 10x ligation buffer (NEB), 100 *μ*l of 10% Triton-X-100, 10 *μ*l of 10 mg/ml BSA, 6.5 *μ*l of T4 DNA ligase (NEB, M0202L), 283.5 *μ*l water) were added to each sample and incubated overnight at 16°C and 800 rpm shaking.

#### Cross-link reversal and DNA purification

Nuclei were pelleted by centrifugation for 10 min, 600 g at 4°C and sample volume was reduced to a total of 200 *μ*l. 230 *μ*l of 10 mM Tris HCL pH 7.5, 20 *μ*l of Proteinase K (10mg/ml) and 50 *μ*l of 10% SDS were added, mixed by pipetting and incubated 30 min at 55°C. Subsequently, 40 *μ*l of 4 M NaCl were added and samples were incubated overnight at 65°C with 700 rpm shaking. Next, 5 *μ*l of RNAse A (10 mg/ml) were added, followed by incubation at 37°C for 30 min at 700 rpm. 20 *μ*l Proteinase K (10 mg/ml) were added to the sample and incubated at 55°C for 1-2 h at 700 rpm. DNA was purified by phenol-chloroform extraction. Following DNA precipitation, dried DNA pellet was reconstituted in 100 *μ*l 10 mM Tris-HCl pH 7.5.

#### Removing biotin from un-ligated fragments and DNA shearing

5-7 *μ*g of HiC library in a total volume of 100 *μ*l (1x NEB buffer 2.1, 0.025 mM dNTPs, 0.12 U/ *μ*l T4 DNA polymerase (NEB, M0203) was incubated at 20°C for 4 h to remove biotin from unligated ends. Reaction was stopped by adding EDTA to a final concentration of 10 mM and heat inactivation for 20 min at 75°C. DNA was sheared, using Covaris M220 sonicator with the following setup: 130 *μ*l sample volume, Peak Incident Power (W): 50, Duty Factor: 20%, Cycles per Burst: 200, Treatment Time (s): 65, cooling at 7°C. Samples were subsequently size selected for fragments between 150 and 600 bp using AMPure XP beads (Agencourt, A63881) as follows: 0.575x volume of AMPure beads were added to the sample, mixed by pipetting, and incubated for 10 min at room temperature. Beads were separated on a magnet, and clear supernatant was transferred to a fresh tube. 0.395x volume of fresh AMPure beads were added to the supernatant, mixed, and incubated for 10 min at room temperature. Beads were separated on a magnet, and clear supernatant was discarded. Beads were washed twice with 70% EtOH, air dried for 5 min and DNA was eluted in 300 *μ*l water.

#### Biotin pull down

Biotin-labelled DNA was bound to Dynabeads My One C1 Streptavidin beads, using 5 *μ*l of beads per 1 *μ*g DNA and following manufacturer’s instructions. Beads were washed twice with 1x tween-washing-buffer (5 mM Tris HCl pH 7.5, 0.5 mM EDTA, 1 M NaCl, 0.05% Tween 20) and finally resuspended in 1x sample volume 2x binding buffer (10 mM Tris HCl pH 7.5, 1 mM EDTA, 2 M NaCl). Beads were mixed with the DNA sample and incubated for 20 min at room temperature while rotating. Beads were separated on a magnet, twice washed with 1x tween-washing-buffer at 55°C and 700 rpm shaking for 2 min. Reclaimed beads were resuspended in 50 *μ*l water.

#### Sequencing library preparation

To repair DNA ends, DNA-bound beads were incubated in 100 *μ*l end-repair mix containing 1x T4 Ligase Buffer (NEB), 0.5 mM dNTP mix, 0.5 U/ *μ*l T4 Polynucleotide Kinase (NEB, M0201), 0.12 U/ *μ*l T4 DNA Polymerase (NEB, M0203) and 0.05 U/ *μ*l Klenow (NEB, M0210). Samples were incubated for 30 min at 20°C. Beads were separated on a magnet, twice washed with 1x tween-washing-buffer at 55°C and 700 rpm shaking for 2 min. Reclaimed beads were resuspended in 50 *μ*l water. Next, dA-tail was added by incubating DNA-bound beads in 100 *μ*l A-tailing mix, containing 1x NEB buffer, 0.5 *μ*M dATP, and 0.25 U/ *μ*l Klenow, exo-(NEB, M0212). Samples were incubated for 30 min at 37°C. Beads were separated on a magnet, twice washed with 1x tween-washing-buffer at 55°C and 700 rpm shaking for 2 min. Reclaimed beads were resuspended in 20 *μ*l water. Subsequently, samples were indexed by ligating TruSeq Illumina adaptors by incubating DNA-bound beads in 50 *μ*l adapter ligation mix, containing 1x T4 Ligation buffer, 5% PEG-4000, 0.3 U/ *μ*l T4 DNA Ligase (ThermoFisher, EL0011), 1.5 *μ*l TruSeq index adapter. The reaction was incubated at 22°C for 2 hours with occasionally mixing. Beads were separated on a magnet, twice washed with 1x tween-washing-buffer at 55°C and 700 rpm shaking for 2 min. Reclaimed beads were resuspended in 50 *μ*l water. Final library for paired-end sequencing was prepared using NEBNext High-Fidelity 2X PCR Master Mix (NEB). PCR reaction: 50 *μ*l reaction, containing 1x NEBNext High-Fidelity PCR Master Mix, 0.3 *μ*M TruSeq Primer 1.0 (P5) and TruSeq Primer 2.0 (P7), 3 *μ*l DNA-bound beads. PCR cycler setup: 1. 98°C for 60 seconds, 2. 98°C for 10 seconds, 3. 65°C for 30 seconds, 4. 72°C for 30 seconds, 5. Go to step 2 for up to 10 cycles, 6. 72°C for 5 min. Optimal cycle number was determined for each sample by analysing a 5 *μ*l aliquot on an agarose gel after 4, 6, 8, 10 and 12 cycles. For each sample, at least 8 independent PCR reactions were performed to maintain initial library complexity and then pooled for AMPure beads purification. 1.2x volume of AMPure beads were added to the sample, mixed by pipetting, and incubated for 10 min at room temperature. Beads were separated on a magnet, and clear supernatant was discarded. Beads were washed twice with 70% EtOH, and air dried for 5 min. DNA was eluted in 50 *μ*l water. Libraries were multiplexed and sequenced using DNBseq technology to produce 50 bp paired-end reads and approximately 400 million raw sequencing read pairs for each genotype.

### HiC data analyses

#### Mapping, filtering, normalization and visualization

HiC paired-end reads were mapped to the zebrafish genome assembly GRCz10 (danRer10) using BWA^68^. Reads from biological replicates were pooled before mapping. Then, ligation events (HiC pairs) were detected and sorted, and PCR duplicates were removed, using the pairtools package (https://github.com/mirnylab/pairtools). Unligated and self-ligated events (dangling and extra-dangling ends, respectively) were filtered out by removing contacts mapping to the same or adjacent restriction fragments. The resulting filtered pairs file was converted to a tsv file that was used as input for Juicer Tools Pre^69^, which generated multiresolution hic files. HiC matrices at 10 and 500 Kb resolution, normalized with the Knight-Ruiz (KR) method^70^, were extracted for downstream analysis using the FAN-C toolkit^71^. Visualization of normalized HiC matrices and other values described below, such as insulation scores, TAD boundaries, aggregate TAD and loop analysis, Pearson’s correlation matrices and eigenvectors, were calculated and visualized using FAN-C.

#### TADs, chromatin loops and compartmentalization

TAD boundaries were called using the insulation score method, as previously described^40^. Insulation scores were calculated for 10-Kb binned HiC matrices using FAN-C^71^. Briefly, the average number of interactions of each bin were calculated in 500-Kb square sliding windows (50 x 50 bins); then, these values were normalized as the log_2_ ratio of each bin’s value and the mean of all bins to obtain the insulation score for each bin; next, minima along the insulation score vector were calculated using a delta vector of +/−100 Kb (+/−10 bins) around the central bin; finally, boundaries with scores lower than 0.5 were filtered out. The genomic regions located between adjacent boundaries were considered as TADs.

For determination of A and B compartments, 500-Kb binned HiC matrices were used. Pearson’s correlation matrices were calculated as previously described^72^, using FAN-C^71^. A/B compartments and their strength were determined using the 2^nd^ eigenvector, since the 1^st^ eigenvector corresponded with chromosome arms in our system, and the genome GC content. A/B domains were defined as consecutive regions with the same eigenvector sign. A/B enrichment profiles were calculated by dividing bins in fifty percentiles according to their 2^nd^ eigenvector values and plotting their average observed/expected contact values.

Chromatin loops were called using HICCUPS^10^, with standard parameters. Briefly, the multiresolution hic file was used as input for the CPU version of HICCUPS, which run using 5, 10 and 25-Kb resolution KR-normalized matrices. The maximum permitted FDR value was 0.1 for the three resolutions; the peak widths were 4, 2 and 1 bin for 5, 10 and 25-Kb resolutions, respectively; and the window widths to define the local neighborhoods used as background were 7, 5 and 3 bins, respectively. The thresholds for merging loop lists from different resolutions were the following: maximum sum of FDR values of 0.02 for the horizontal, vertical, donut and lower-left neighborhoods; minimum enrichment of 1.5 for the horizontal and vertical neighborhoods; minimum enrichment of 1.75 for the donut and bottom-left neighborhoods; minimum enrichment of 2 for either the donut or the bottom-left neighborhoods. The distances used to merge the nearby pixels to a centroid were 20, 20 and 50-Kb for 5, 10 and 25-Kb resolutions, respectively. CTCF-bound and chromatin loops were considered when at least one of the loop anchors overlapped with a CTCF ChIP-seq peak.

### UMI-4C

UMI-4C library preparation was performed as previously described^73^ with modifications in 3C library preparation and minor modification in sequencing library preparation. Experiments were performed in singletons in wild-type and *ctcf*^−/−^ mutant embryos at 48 hpf, using one to three million cells as input material. Embryo fixation, nuclei extraction, chromatin digestion, biotin fill-in, proximity ligation, cross-link reversal, and DNA purification were performed following above experimental procedure for HiC. The following procedure were specific for UMI-4C.

#### DNA shearing

5-7 *μ*g of purified DNA was sheared with Covaris M220 sonicator with the following setup: 130 *μ*l sample volume, Peak Incident Power (W): 50, Duty Factor: 10%, Cycles per Burst: 200, Treatment Time (s): 70, cooling at 7°C. Samples were then purified using AMPure XP beads (Agencourt, A63881) as follows: 2.0x volume of AMPure beads were added to the sample, mixed by pipetting, and incubated for 10 min at room temperature. Beads were separated on a magnet, and clear supernatant was discarded. Beads were washed twice with 70% EtOH, and air dried for 5 min. DNA was eluted in 300 *μ*l water.

#### Biotin pull down

Biotin-labelled DNA was bound to Dynabeads My One C1 Streptavidin beads, using 5 *μ*l of beads per 1 *μ*g DNA and following manufacturer’s instructions. Beads were washed twice with 1x tween-washing-buffer (5 mM Tris HCl pH 7.5, 0.5 mM EDTA, 1 M NaCl, 0.05% Tween 20) and finally resuspended in 1x sample volume 2x binding buffer (10 mM Tris HCl pH 7.5, 1 mM EDTA, 2 M NaCl). Beads were mixed with the DNA sample and incubated for 20 min at room temperature while rotating. Beads were separated on a magnet, twice washed with 1x tween-washing-buffer at 55°C and 700 rpm shaking for 2 min. Reclaimed beads were resuspended in 50 *μ*l water.

#### Sequencing library preparation

500 ng of DNA attached to beads were end-repaired by incubating in 100 *μ*l end-repair mix (1x T4 Ligase Buffer (NEB), 0.5 mM dNTP mix, 0.12 U/ *μ*l T4 DNA Polymerase (NEB, M0203) and 0.05 U/ *μ*l Klenow (NEB, M0210)) for 30 min at 20°C. Beads were separated on a magnet, twice washed with 1x tween-washing-buffer at 55°C and 700 rpm shaking for 2 min. Reclaimed beads were resuspended in 50 *μ*l water. Next, DNA-bound beads were incubated for 30 min at 37°C in 100 *μ*l A-tailing mix (1x NEB buffer, 0.5 *μ*M dATP, and 0.25 U/ *μ*l Klenow, exo-(NEB, M0212)). The enzyme was heat inactivated at 75°C for 20 min. For 5’ dephosphorylation of DNA ends, 2 *μ*l of Alkaline Phosphatase, Calf Intestinal (NEB, M0290) was added and samples were incubated at 37°C for 1 hour and with occasionally mixing. Beads were separated on a magnet, twice washed with 1x tween-washing-buffer at 55°C and 700 rpm shaking for 2 min. Reclaimed beads were resuspended in 20 *μ*l water. Next, samples were indexed by ligating TruSeq Illumina adaptors by incubating DNA-bound beads in 50 *μ*l adapter ligation mix (1x T4 Ligation buffer, 5% PEG-4000, 0.3 U/ *μ*l T4 DNA Ligase (ThermoFisher, EL0011), 1.5 *μ*l TruSeq index adapter). The reaction was incubated at 22°C for 2 hours with occasionally mixing. Sample volume was increased with water to a total 100 *μ*l and incubated at 96°C for 5 min to denature DNA and remove non-ligated strand from adapter. Sample were placed on ice and beads were separated on a magnet, twice washed with 1x tween-washing-buffer at 55°C and 700 rpm shaking for 2 min. Reclaimed beads were resuspended in 20 *μ*l water. Final library for paired-end sequencing was prepared using NEBNext High-Fidelity 2X PCR Master Mix (NEB) and a nested PCR approach as described in Schwartzman et al. 2016. Individual viewpoints are defined by US (upstream) and DS (downstream) primers within the DpnII fragment of interest (Extended Data Table 1). US and DS primers were designed with melting temperature of 58°C. DS primers were designed between 5-15 bp from the interrogated DpnII restriction site and containing P5 sequence at their 5’ end. US primers were designed within a region of up to 100 bp of interrogated DpnII restriction site and with only minimal overlap with DS primers. Up to 14 US and DS primers were pooled for multiplex PCR reaction, respectively. First PCR reaction: 50 *μ*l reaction, containing 1x NEBNext High-Fidelity 2X PCR Master Mix, 0.3 *μ*M US primer mix (each) and 0.3 *μ*M TruSeq Primer 2.0 (P7), 200 ng DNA-bound on beads. PCR cycler setup: 1. 98°C for 30 seconds, 2. 98°C for 10 seconds, 3. 58°C for 30 seconds, 4. 72°C for 60 seconds, 5. Go to step 2 for 18 cycles in total, 6. 72°C for 5 min. For each sample two PCR reactions were performed and then pooled for AMPure beads purification. 1.2x volume of AMPure beads were added to the sample, mixed by pipetting, and incubated for 10 min at room temperature. Beads were separated on a magnet, and clear supernatant was discarded. Beads were washed twice with 70% EtOH, air dried, and DNA was eluted in 30 *μ*l water. Second PCR reaction: 50 *μ*l reaction, containing 1x NEBNext High-Fidelity 2X PCR Master Mix, 0.3 *μ*M DS primer mix (each) and 0.3 *μ*M TruSeq Primer 2.0 (P7), 100 ng DNA from first PCR. PCR cycler setup: Corresponded to setup of first PCR but with 15 cycles. For each sample 3-5 PCR reactions were performed and then pooled for size selection for fragments between 200 and 700 bp, using AMPure beads. 0.575x volume of AMPure beads were added to the sample, mixed by pipetting, and incubated for 10 min at room temperature. Beads were separated on a magnet, and clear supernatant was transferred to a fresh tube. 0.3x volume of fresh AMPure beads were added to the supernatant, mixed, and incubated for 10 min at room temperature. Beads were separated on a magnet, and clear supernatant was discarded. Beads were washed twice with 70% EtOH, and air dried for 5 min. DNA was eluted in 300 *μ*l water. Libraries were multiplexed and sequenced using DNBseq technology to produce 50 bp paired-end reads and approximately 1-5 million raw sequencing read pairs for each viewpoint and genotype.

For the UMI-4C data analysis, raw fastq files were processed using the R package umi4cpackage (https://bitbucket.org/tanaylab/umi4cpackage). Contact profiles and domainograms were generated using the default parameters and a minimum win_cov of 10.

### Statistical analyses

For comparison of insulation scores, TAD sizes, loop ranges and expression fold-changes among datasets, two-tailed Wilcoxon’s rank sum tests were used. In Fig. 3f and Extended data Fig. 7, box plots represent: center line, median; box limits, upper and lower quartiles; whiskers, 1.5x interquartile range; notches, 95% confidence interval of the median. Other boxplots represent the same parameters but do not include notches. Statistical significance of contingency tables was assessed using the Fisher’s exact test.

## Data availability

HiC, ChIPmentation, RNA-seq, ATAC-seq and UMI-4C data generated in this study are available through the Gene Expression Omnibus (GEO) accession number GSE156099 [https://www.ncbi.nlm.nih.gov/geo/query/acc.cgi?acc=GSE156099].

## Code availability

Custom code used in this study is available at the Gitlab repository:] https://gitlab.com/rdacemel/hic_ctcf-null.

## Acknowledgements

We thank C. Paliou for critical reading of the manuscript; C. Bolt and L. Delisle from the Duboule lab for technical advice with the UMI-4C protocol; F. Rencillas-Targa for providing the zebrafish-specific CTCF antibody; the CABD Fish and Microscopy Facilities for technical assistance; and C3UPO for the HPC support. JLG-S received funding from the ERC (Grant Agreement No. 740041), the Spanish Ministerio de Economía y Competitividad (Grant No. BFU2016-74961-P) and the institutional grant Unidad de Excelencia María de Maeztu (MDM-2016-0687). JT was funded by a 2019 Leonardo Grant for Researchers and Cultural Creators, BBVA Foundation. MF was funded by the European Union’s Horizon 2020 research and innovation programme under the Marie Skłodowska-Curie grant agreement [#800396] and a Juan de la Cierva-Formación fellow from the Spanish Ministry of Science and Innovation (FJC2018-038233-I).

## Author contributions

MF, JMS-P and JLG-S conceived and designed the project; EC-M, MF, AN and JMS-P performed the experiments; JMS-P, MF, RDA and JJT analyzed the data; MF, JMS-P and JLG-S wrote the manuscript.

**Correspondence and requests for materials** should be addressed to JMS-P or JLG-S.

## EXTENDED DATA

**Extended Data Table 1.**
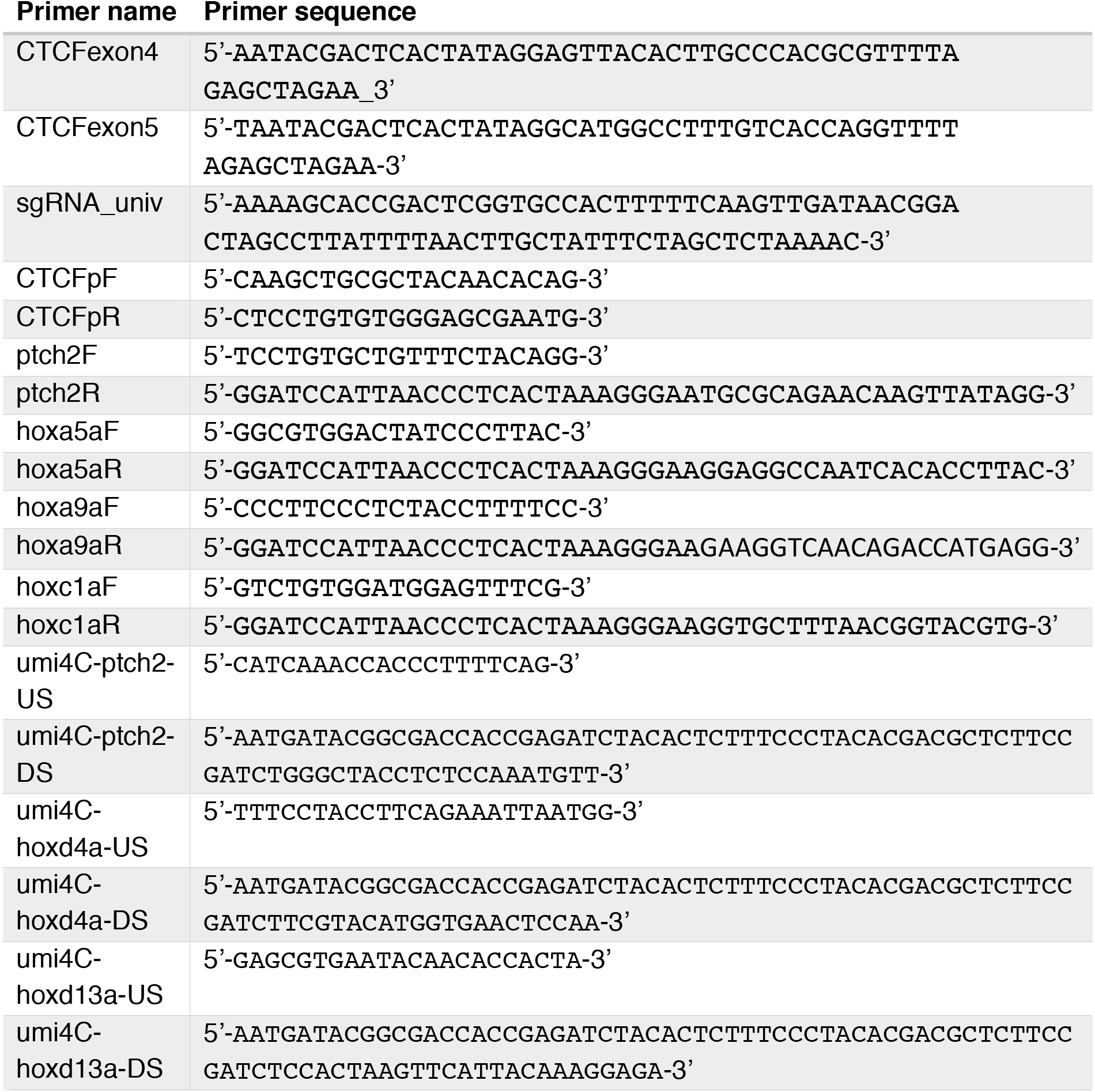
List of primers used in this study

---

**Extended Data Figure 1.**
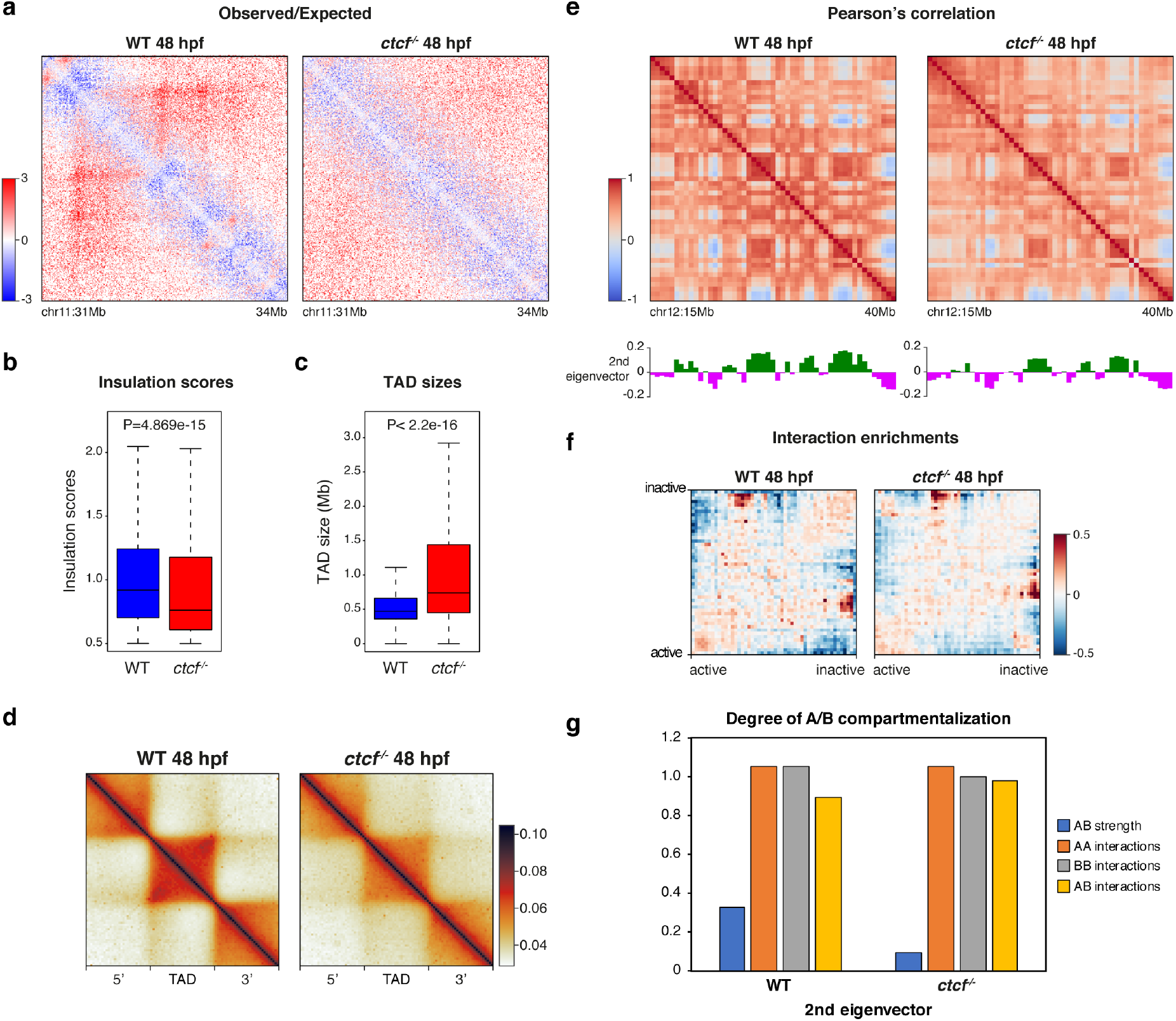
Chromatin structure in zebrafish embryos requires CTCF. **a,** HiC observed/expected contact maps at 10 Kb resolution in WT and *ctcf*^−/−^ zebrafish embryos at 48 hpf. The 3-Mb genomic region shown in Figure 1c is plotted. **b-c,** Box plots showing the insulation scores of the TAD boundaries (b) and the TAD sizes (c) in WT and *ctcf*^−/−^ embryos at 48 hpf. Statistical significance was assessed using the Wilcoxon’s rank sum test. **d,** Aggregate analysis of normalized HiC signal in WT and *ctcf*^−/−^ embryos at 48 hpf for the 2,438 TADs called in WT embryos, rescaled and surrounded by windows of the same size. **e,** Pearson’s correlation matrices from HiC data at 500 Kb resolution in WT and *ctcf*^−/−^ embryos at 48 hpf. A 25-Mb genomic region is plotted, aligned with the 2^nd^ eigenvector demarcating A and B compartments. **f,** Saddle plots showing the genome-wide interaction enrichments between active and inactive genomic regions from HiC data in WT and *ctcf*^−/−^ embryos at 48 hpf. **g,** Bar plots showing the degree of compartmentalization in WT and *ctcf*^−/−^ embryos at 48 hpf. AB strength and quantification of AA, BB and AB interactions based in the 2^nd^ eigenvector, are plotted.

**Extended Data Figure 2.**
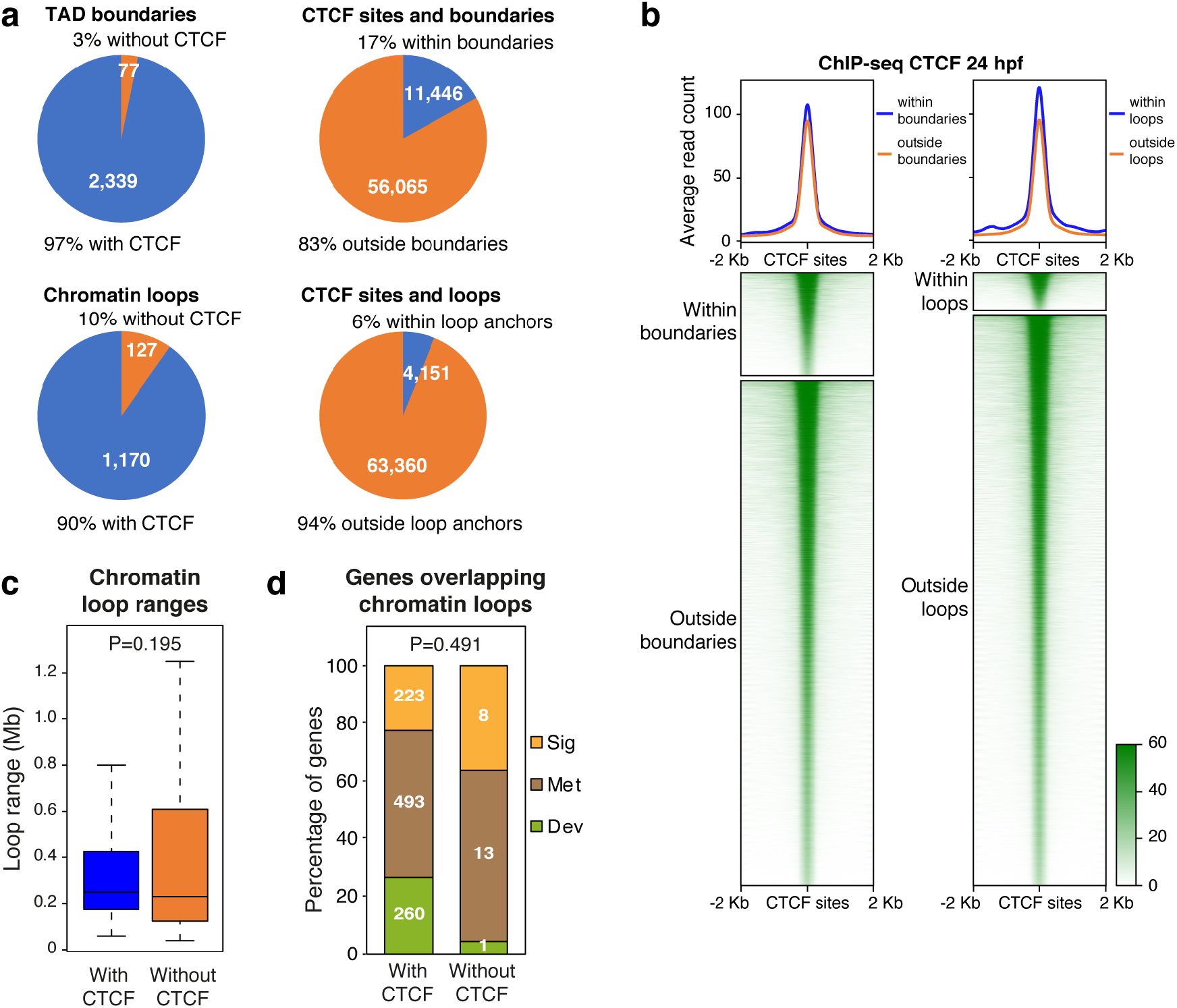
CTCF is bound to TAD boundaries and chromatin loops in zebrafish embryos. **a,** Pie charts showing the percentage of TADs or chromatin loops overlapping with CTCF sites (left) and the percentage of CTCF sites overlapping with TADs or chromatin loops (right). **b,** Heatmaps and average profiles of CTCF ChIP-seq signal at CTCF sites overlapping or not with TADs or chromatin loops. **c,** Box plots showing the distance between loop anchors (loop ranges) for the chromatin loops overlapping or not with CTCF sites at least in one of their anchors. Statistical significance was assessed using the Wilcoxon’s rank sum test. **d,** Proportion of genes annotated to the GO terms “Signaling”, “Metabolic process” and “Developmental process” for genes overlapping with chromatin loop anchors, with or without CTCF binding. Statistical significance was assessed using the Fisher’s exact test.

**Extended Data Figure 3.**
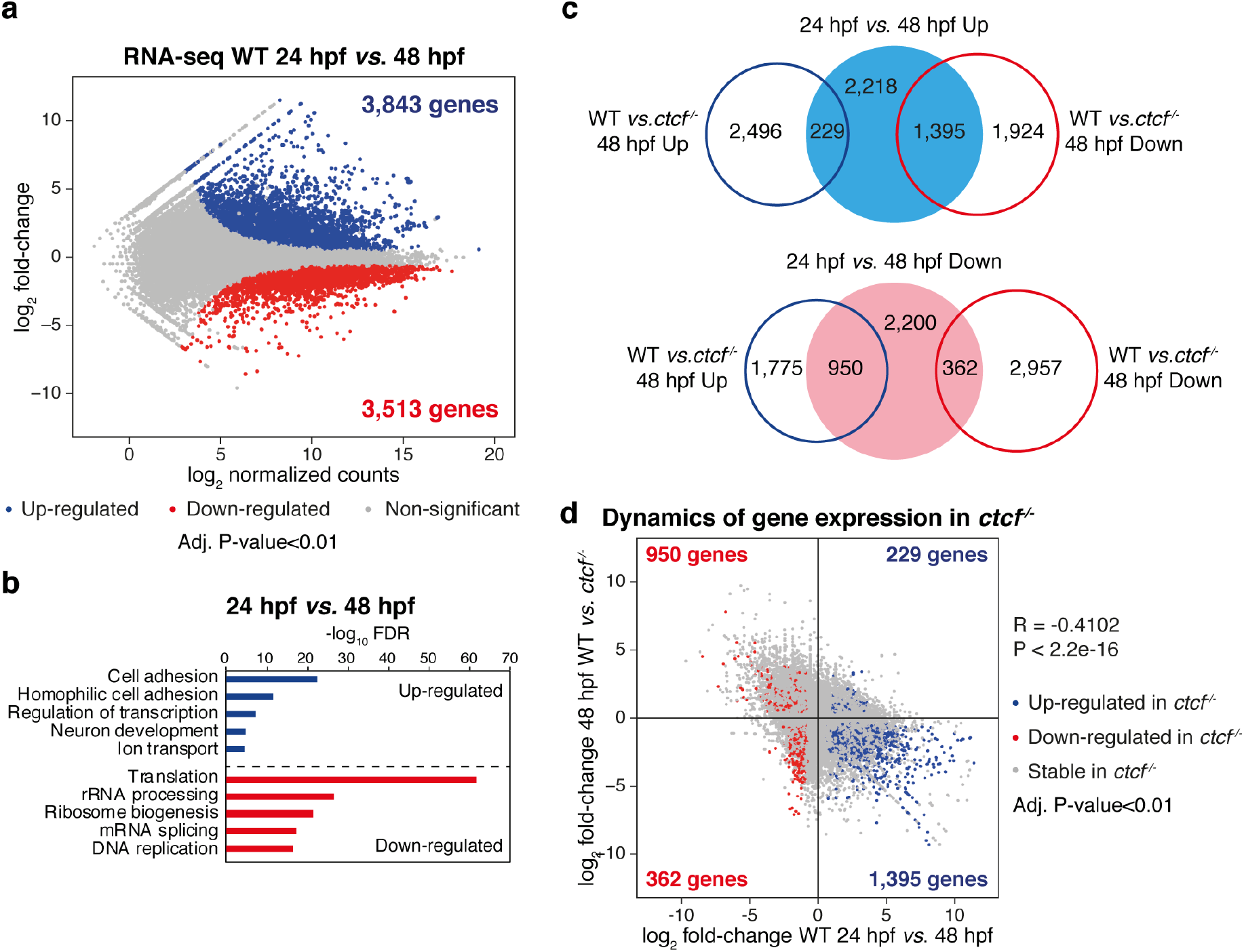
CTCF is required for dynamic expression changes during development. **a,** Differential analysis of gene expression in WT embryos between 24 and 48 hpf from RNA-seq data (n = 2 biological replicates per condition). The log_2_ normalized read counts of 24-hpf transcripts versus the log_2_ fold-change of expression are plotted. Transcripts showing a statistically significant differential expression (adjusted P-value < 0.01) are highlighted in blue (up-regulated) or red (down-regulated). The number of genes that correspond to the up- and down-regulated transcripts are shown inside the boxes. **b,** GO enrichment analyses of biological processes for the up- and down-regulated genes in WT embryos from 24 to 48 hpf. Terms showing an FDR < 0.05 are considered as enriched. **c,** Venn diagrams showing the overlap between the genes up- and down-regulated in WT embryos from 24 to 48 hpf and the genes up- and down-regulated in *ctcf*^−/−^ embryos at 48 hpf (see Fig. 2b). **d,** Scatter plots showing the correlation between the expression fold change of all transcripts in WT embryos from 24 to 48 hpf, and their expression fold change in *ctcf*^−/−^ embryos at 48 hpf. Up- and down-regulated transcripts in *ctcf*^−/−^ embryos are highlighted in blue or red, respectively.

**Extended Data Figure 4.**
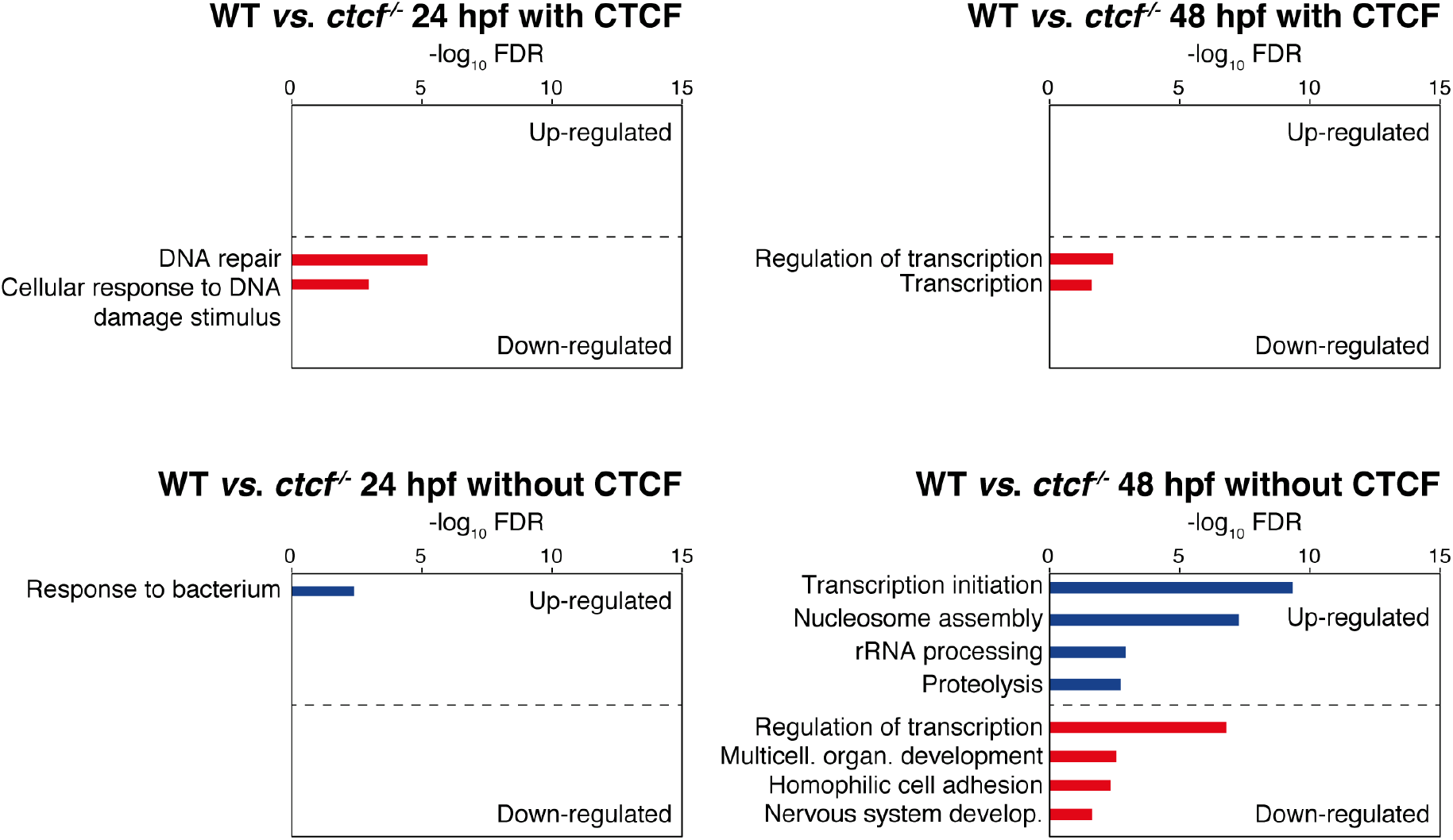
CTCF absence leads to down-regulation of developmental genes. GO enrichment analyses of biological processes for the up- and down-regulated genes in *ctcf*^−/−^ embryos at 24 (left) and 48 hpf (right), distinguishing between those genes with (top) or without (bottom) CTCF binding at their TSS. Terms showing a false discovery rate (FDR) < 0.05 are considered as enriched.

**Extended Data Figure 5.**
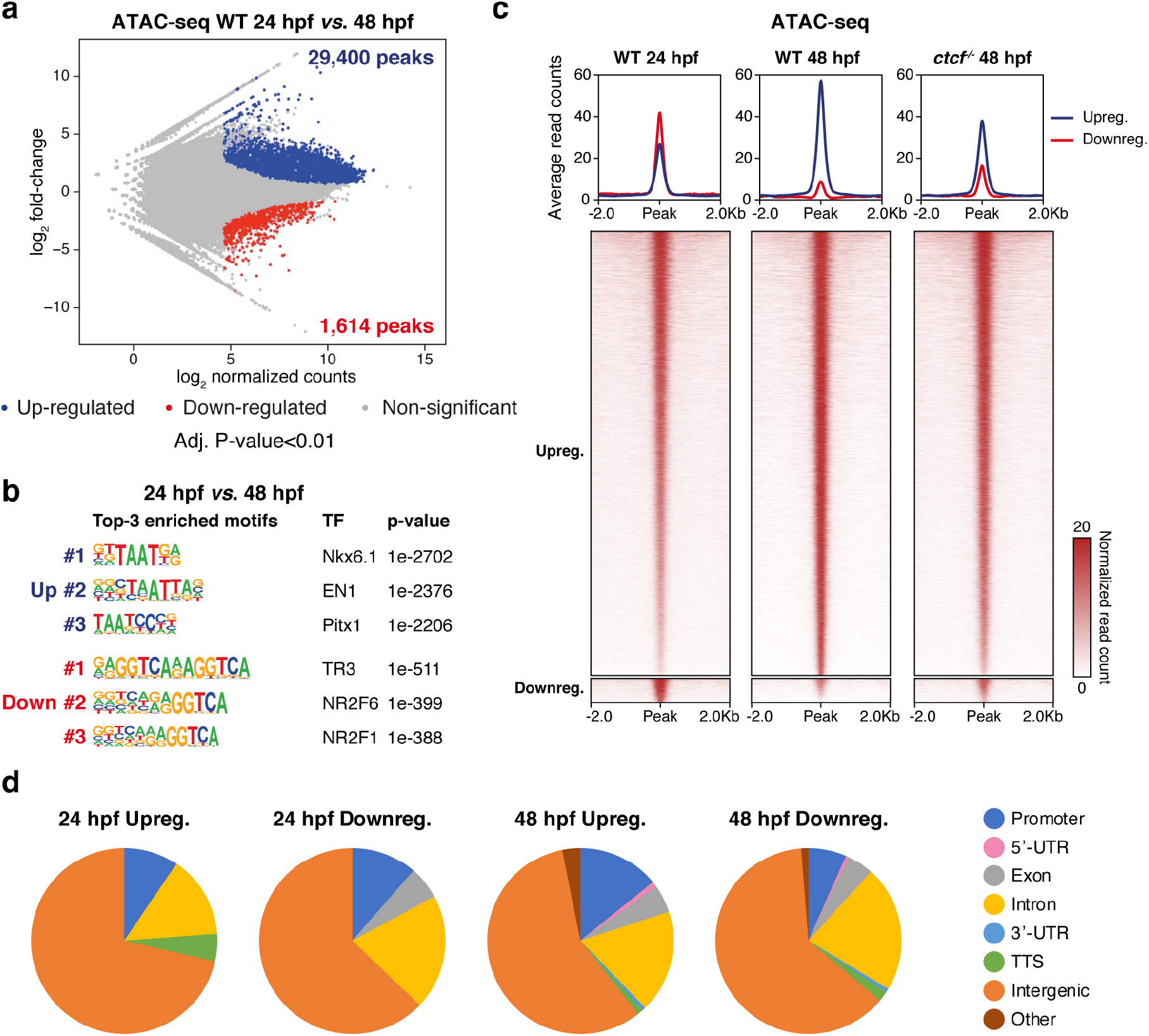
The dynamics of the chromatin accessibility landscape requires CTCF. **a,** Differential analysis of chromatin accessibility in WT embryos between 24 and 48 hpf from ATAC-seq data (n = 2 biological replicates per condition). The log_2_ normalized read counts of 24-hpf ATAC peaks versus the log_2_ fold-change of accessibility are plotted. Regions showing a statistically significant differential accessibility (adjusted P-value < 0.01) are highlighted in blue (up-regulated) or red (down-regulated). The number of peaks that correspond to the up- and down-regulated sites are shown inside the boxes. **b,** Motif enrichment analyses for the up- and down-regulated ATAC peaks in WT embryos from 24 to 48 hpf. The 3 motifs with the lowest p-values are shown for each case. **c,** Heatmaps and average profiles plotting normalized ATAC-seq signal in WT embryos at 24 and 48 hpf and in *ctcf*^−/−^ embryos at 48 hpf for the up- and down-regulated peaks from (a). **d,** Pie charts showing the annotation to different genomic features of ATAC-peaks up- or down-regulated in *ctcf*^−/−^embryos at 24 or 48 hpf (see Fig. 3a-b).

**Extended Data Figure 6.**
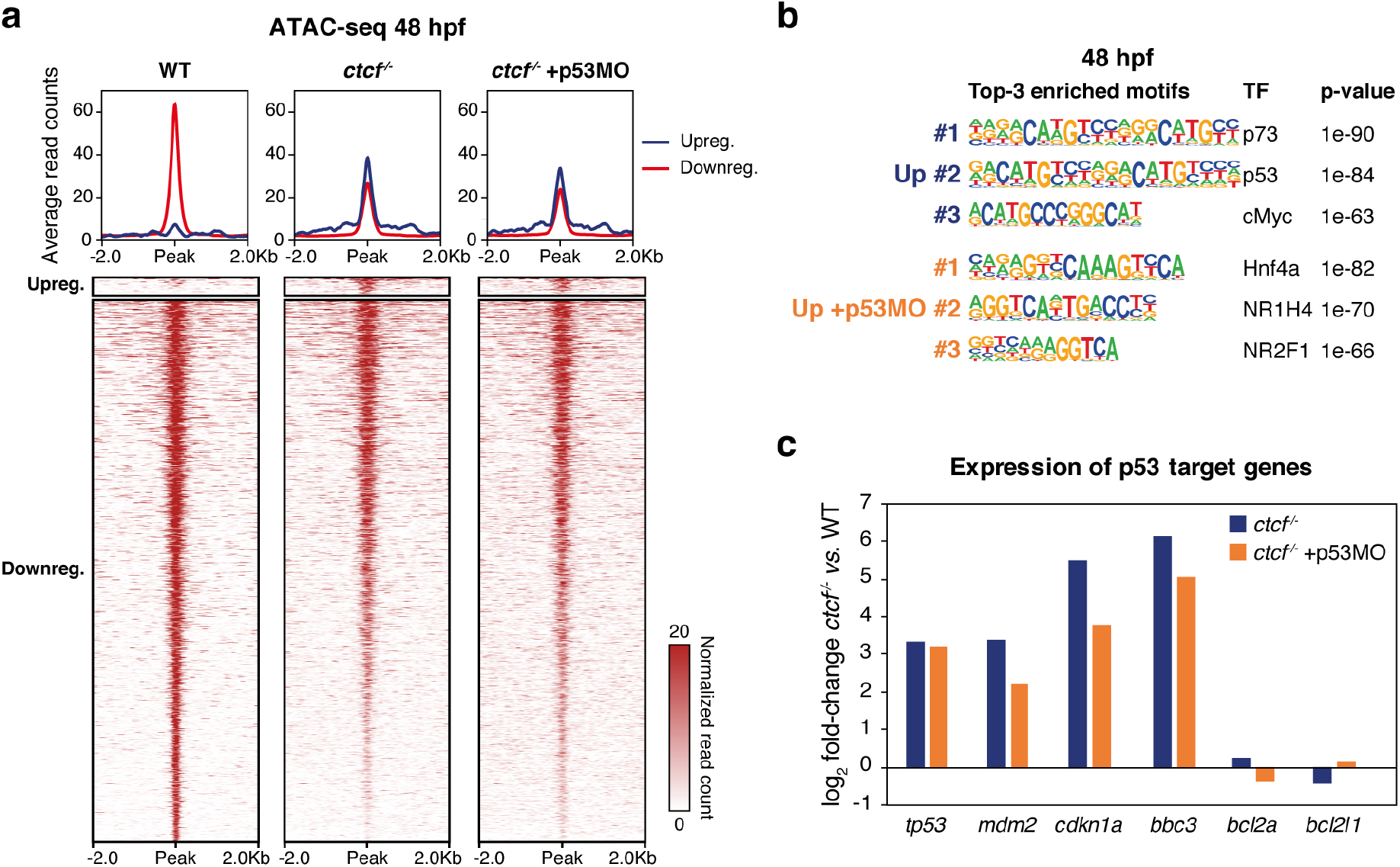
The p53 pro-apoptotic response in the absence of CTCF does not suppress defects in chromatin accessibility. **a,** Heatmaps and average profiles plotting normalized ATAC-seq signal in WT, control *ctcf*^−/−^ and p53 morpholino (p53MO)-injected *ctcf*^−/−^ embryos at 48 hpf for the up- and down-regulated peaks in control *ctcf*^−/−^ embryos (see Fig. 3b). **b,** Motif enrichment analyses for the up-regulated ATAC peaks in control *ctcf*^−/−^ and p53MO-injected *ctcf*^−/−^ embryos at 48 hpf. The 3 motifs with the lowest p-values are shown for each case. **c,** Gene expression fold change from RNA-seq data of the *tp53* gene and the p53 target genes *mdm2*, *cdkn1a*, *bbc3*, *bcl2a* and *bdl2l1*, in control *ctcf*^−/−^ and p53MO-injected *ctcf*^−/−^ embryos at 48 hpf.

**Extended Data Figure 7.**
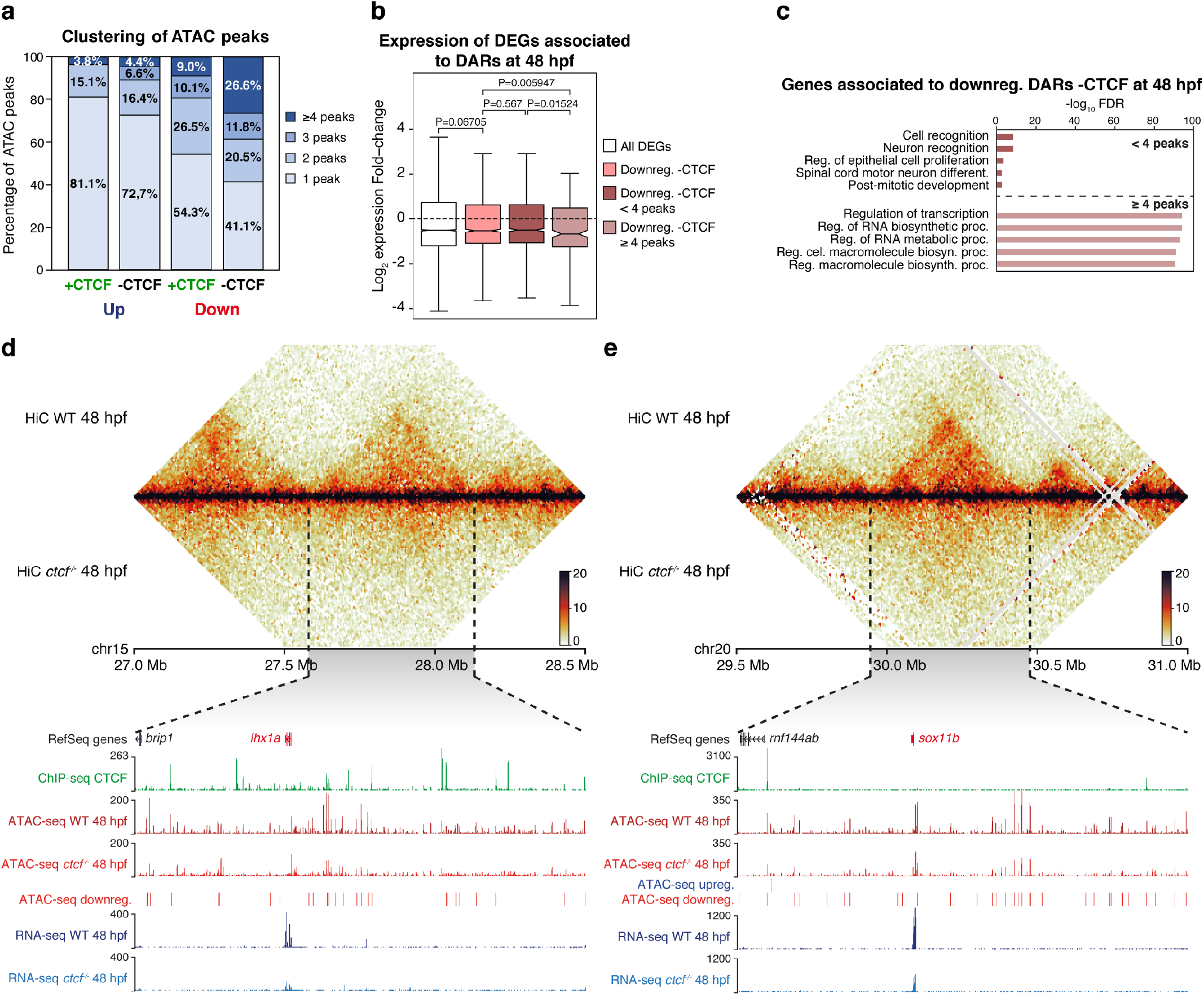
CTCF loss reduces accessibility at clustered *cis*-regulatory elements around developmental genes. **a,** Bar plots showing the level of clustering of the up- and down-regulated ATAC-seq peaks in *ctcf*^−/−^ embryos at 48 hpf, with or without CTCF binding. Peaks were considered to be clustered when located less than 30 Kb from each other. **b,** Box plots showing the expression fold-change in *ctcf*^−/−^ embryos at 48 hpf of all DEGs or only those associated with down-regulated DARs not overlapping with CTCF sites, grouped in less or more than 4 peaks per cluster. Center line, median; box limits, upper and lower quartiles; whiskers, 1.5x interquartile range; notches, 95% confidence interval of the median Statistical significance was assessed using the Wilcoxon’s rank sum test. **c,** GO enrichment analyses of biological processes for the genes associated with the down-regulated DARs in *ctcf*^−/−^ embryos at 48 hpf not overlapping with CTCF sites, grouped in less or more than 4 peaks per cluster. Top-5 terms showing an FDR < 0.05 are considered as enriched. **d,** Top, heatmaps showing HiC signal in WT and *ctcf*^−/−^ embryos at 48 hpf in a 1.5-Mb region of chromosomes 15 (left) or 20 (right). Bottom, zoom within the *lhx1a* TAD (left) or the *sox11b* TAD (right) showing UCSC Genome Browser tracks with CTCF ChIP-seq, ATAC-seq at 48 hpf in WT and *ctcf*^−/−^ embryos, ATAC-seq up- or down-regulated peaks and RNA-seq at 48 hpf in WT and *ctcf*^−/−^ embryos. The down-regulated genes are shown in red.

**Extended Data Figure 8.**
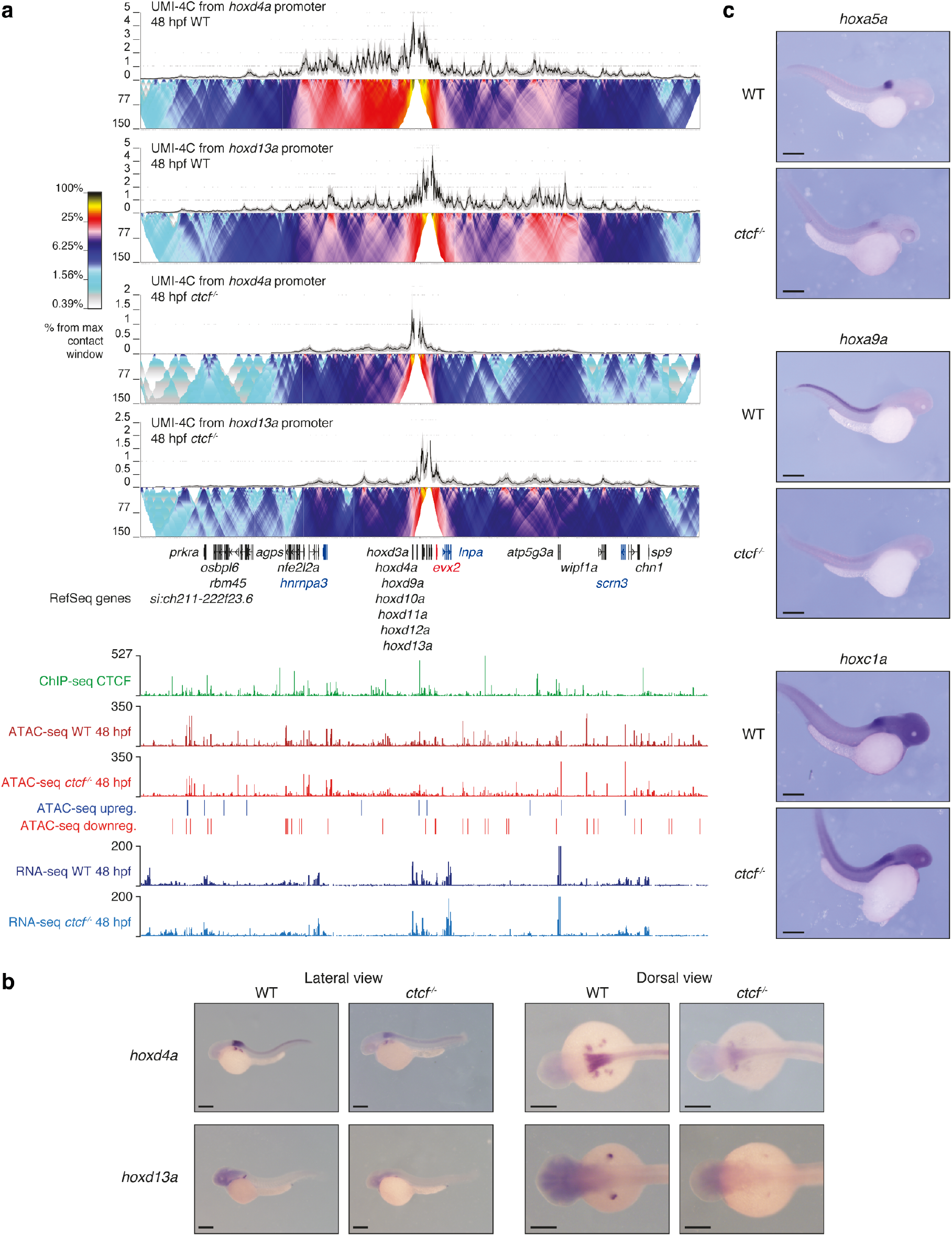
CTCF is required for the establishment of regulatory landscapes at the HoxD locus and *hox* gene expression. **a,** Top, UMI-4C assays in WT and *ctcf*^−/−^ embryos at 48 hpf using the *hoxd4a* and *hoxd13a* gene promoters as viewpoints. Black lines and grey shadows represent the average normalized UMI counts and their standard deviation, respectively. Domainograms below UMI counts represent contact frequency between pairs of genomic regions. Bottom, UCSC Genome Browser tracks with CTCF ChIP-seq, ATAC-seq at 48 hpf in WT and *ctcf*^−/−^ embryos, ATAC-seq up- and down-regulated peaks and RNA-seq at 48 hpf in WT and *ctcf*^−/−^ embryos. Up- and down-regulated genes are shown in blue and red, respectively. **b,** Whole-mount *in situ* hybridization of the *hoxd4a* and *hoxd13a* genes in WT and *ctcf*^−/−^ embryos at 48 hpf. Left, lateral view; right, dorsal view. Anterior is to the left and scale bars represent 500 *μ*m. **c,** Whole-mount *in situ* hybridization of the *hoxa5a*, *hoxa9a* and *hoxc1a* genes in WT and *ctcf*^−/−^ embryos at 48 hpf. Left, Anterior is to the right and scale bars represent 500 *μ*m.

## References

1 Bell, A. C., West, A. G. & Felsenfeld, G. The protein CTCF is required for the enhancer blocking activity of vertebrate insulators. Cell 98, 387–396, doi:10.1016/s0092-8674(00)81967-4 (1999).

2 Filippova, G. N. et al. An exceptionally conserved transcriptional repressor, CTCF, employs different combinations of zinc fingers to bind diverged promoter sequences of avian and mammalian c-myc oncogenes. Mol Cell Biol 16, 2802–2813, doi:10.1128/mcb.16.6.2802 (1996).

3 Merkenschlager, M. & Nora, E. P. CTCF and Cohesin in Genome Folding and Transcriptional Gene Regulation. Annu Rev Genomics Hum Genet 17, 17–43, doi:10.1146/annurev-genom-083115-022339 (2016).

4 Phillips, J. E. & Corces, V. G. CTCF: master weaver of the genome. Cell 137, 1194–1211, doi:10.1016/j.cell.2009.06.001 (2009).

5 Nora, E. P. et al. Targeted Degradation of CTCF Decouples Local Insulation of Chromosome Domains from Genomic Compartmentalization. Cell 169, 930–944 e922, doi:10.1016/j.cell.2017.05.004 (2017).

6 Dixon, J. R. et al. Topological domains in mammalian genomes identified by analysis of chromatin interactions. Nature 485, 376–380, doi:10.1038/nature11082 (2012).

7 Hou, C., Li, L., Qin, Z. S. & Corces, V. G. Gene density, transcription, and insulators contribute to the partition of the Drosophila genome into physical domains. Mol Cell 48, 471–484, doi:10.1016/j.molcel.2012.08.031 (2012).

8 Nora, E. P. et al. Spatial partitioning of the regulatory landscape of the X-inactivation centre. Nature 485, 381–385, doi:10.1038/nature11049 (2012).

9 Sexton, T. et al. Three-dimensional folding and functional organization principles of the Drosophila genome. Cell 148, 458–472, doi:10.1016/j.cell.2012.01.010 (2012).

10 Rao, S. S. et al. A 3D map of the human genome at kilobase resolution reveals principles of chromatin looping. Cell 159, 1665–1680, doi:10.1016/j.cell.2014.11.021 (2014).

11 Splinter, E. et al. CTCF mediates long-range chromatin looping and local histone modification in the beta-globin locus. Genes Dev 20, 2349–2354, doi:10.1101/gad.399506 (2006).

12 Phillips-Cremins, J. E. et al. Architectural protein subclasses shape 3D organization of genomes during lineage commitment. Cell 153, 1281–1295, doi:10.1016/j.cell.2013.04.053 (2013).

13 Heath, H. et al. CTCF regulates cell cycle progression of alphabeta T cells in the thymus. EMBO J 27, 2839–2850, doi:10.1038/emboj.2008.214 (2008).

14 Moore, J. M. et al. Loss of maternal CTCF is associated with peri-implantation lethality of Ctcf null embryos. PLoS One 7, e34915, doi:10.1371/journal.pone.0034915 (2012).

15 Chen, X. et al. Key role for CTCF in establishing chromatin structure in human embryos. Nature 576, 306–310, doi:10.1038/s41586-019-1812-0 (2019).

16 de Laat, W. & Duboule, D. Topology of mammalian developmental enhancers and their regulatory landscapes. Nature 502, 499–506, doi:10.1038/nature12753 (2013).

17 Furlong, E. E. M. & Levine, M. Developmental enhancers and chromosome topology. Science 361, 1341–1345, doi:10.1126/science.aau0320 (2018).

18 Franke, M. & Gomez-Skarmeta, J. L. An evolutionary perspective of regulatory landscape dynamics in development and disease. Curr Opin Cell Biol 55, 24–29, doi:10.1016/j.ceb.2018.06.009 (2018).

19 Bonev, B. & Cavalli, G. Organization and function of the 3D genome. Nat Rev Genet 17, 661–678, doi:10.1038/nrg.2016.112 (2016).

20 Dekker, J. & Mirny, L. The 3D Genome as Moderator of Chromosomal Communication. Cell 164, 1110–1121, doi:10.1016/j.cell.2016.02.007 (2016).

21 Dixon, J. R., Gorkin, D. U. & Ren, B. Chromatin Domains: The Unit of Chromosome Organization. Mol Cell 62, 668–680, doi:10.1016/j.molcel.2016.05.018 (2016).

22 McCord, R. P., Kaplan, N. & Giorgetti, L. Chromosome Conformation Capture and Beyond: Toward an Integrative View of Chromosome Structure and Function. Mol Cell 77, 688–708, doi:10.1016/j.molcel.2019.12.021 (2020).

23 Rowley, M. J. & Corces, V. G. Organizational principles of 3D genome architecture. Nat Rev Genet 19, 789–800, doi:10.1038/s41576-018-0060-8 (2018).

24 Fudenberg, G. et al. Formation of Chromosomal Domains by Loop Extrusion. Cell Rep 15, 2038–2049, doi:10.1016/j.celrep.2016.04.085 (2016).

25 Sanborn, A. L. et al. Chromatin extrusion explains key features of loop and domain formation in wild-type and engineered genomes. Proc Natl Acad Sci U S A 112, E6456–6465, doi:10.1073/pnas.1518552112 (2015).

26 Rao, S. S. P. et al. Cohesin Loss Eliminates All Loop Domains. Cell 171, 305–320 e324, doi:10.1016/j.cell.2017.09.026 (2017).

27 Franke, M. et al. Formation of new chromatin domains determines pathogenicity of genomic duplications. Nature 538, 265–269, doi:10.1038/nature19800 (2016).

28 Laugsch, M. et al. Modeling the Pathological Long-Range Regulatory Effects of Human Structural Variation with Patient-Specific hiPSCs. Cell Stem Cell 24, 736–752 e712, doi:10.1016/j.stem.2019.03.004 (2019).

29 Lupianez, D. G. et al. Disruptions of topological chromatin domains cause pathogenic rewiring of gene-enhancer interactions. Cell 161, 1012–1025, doi:10.1016/j.cell.2015.04.004 (2015).

30 Spielmann, M., Lupianez, D. G. & Mundlos, S. Structural variation in the 3D genome. Nat Rev Genet 19, 453–467, doi:10.1038/s41576-018-0007-0 (2018).

31 Symmons, O. et al. The Shh Topological Domain Facilitates the Action of Remote Enhancers by Reducing the Effects of Genomic Distances. Dev Cell 39, 529–543, doi:10.1016/j.devcel.2016.10.015 (2016).

32 Kubo, N. et al. CTCF Promotes Long-range Enhancer-promoter Interactions and Lineage-specific Gene Expression in Mammalian Cells. bioRxiv, 2020.2003.2021.001693, doi:10.1101/2020.03.21.001693 (2020).

33 Ghavi-Helm, Y. et al. Highly rearranged chromosomes reveal uncoupling between genome topology and gene expression. Nat Genet 51, 1272–1282, doi:10.1038/s41588-019-0462-3 (2019).

34 Williamson, I. et al. Developmentally regulated Shh expression is robust to TAD perturbations. Development 146, doi:10.1242/dev.179523 (2019).

35 Despang, A. et al. Functional dissection of the Sox9-Kcnj2 locus identifies nonessential and instructive roles of TAD architecture. Nat Genet 51, 1263–1271, doi:10.1038/s41588-019-0466-z (2019).

36 Hnisz, D. et al. Activation of proto-oncogenes by disruption of chromosome neighborhoods. Science 351, 1454–1458, doi:10.1126/science.aad9024 (2016).

37 Paliou, C. et al. Preformed chromatin topology assists transcriptional robustness of Shh during limb development. Proc Natl Acad Sci U S A 116, 12390–12399, doi:10.1073/pnas.1900672116 (2019).

38 Vejnar, C. E. et al. Genome wide analysis of 3’ UTR sequence elements and proteins regulating mRNA stability during maternal-to-zygotic transition in zebrafish. Genome Res 29, 1100–1114, doi:10.1101/gr.245159.118 (2019).

39 Kaaij, L. J. T., van der Weide, R. H., Ketting, R. F. & de Wit, E. Systemic Loss and Gain of Chromatin Architecture throughout Zebrafish Development. Cell Rep 24, 1–10 e14, doi:10.1016/j.celrep.2018.06.003 (2018).

40 Crane, E. et al. Condensin-driven remodelling of X chromosome topology during dosage compensation. Nature 523, 240–244, doi:10.1038/nature14450 (2015).

41 Niu, L. et al. Systematic Chromatin Architecture Analysis in *Xenopus tropicalis* Reveals Conserved Three-Dimensional Folding Principles of Vertebrate Genomes. bioRxiv, 2020.2004.2002.021378, doi:10.1101/2020.04.02.021378 (2020).

42 de Wit, E. et al. CTCF Binding Polarity Determines Chromatin Looping. Mol Cell 60, 676–684, doi:10.1016/j.molcel.2015.09.023 (2015).

43 Gomez-Marin, C. et al. Evolutionary comparison reveals that diverging CTCF sites are signatures of ancestral topological associating domains borders. Proc Natl Acad Sci U S A 112, 7542–7547, doi:10.1073/pnas.1505463112 (2015).

44 Thiecke, M. J. et al. Cohesin-dependent and independent mechanisms support chromosomal contacts between promoters and enhancers. bioRxiv, 2020.2002.2010.941989, doi:10.1101/2020.02.10.941989 (2020).

45 Stik, G. et al. CTCF is dispensable for immune cell transdifferentiation but facilitates an acute inflammatory response. Nat Genet, doi:10.1038/s41588-020-0643-0 (2020).

46 Harmston, N. et al. Topologically associating domains are ancient features that coincide with Metazoan clusters of extreme noncoding conservation. Nat Commun 8, 441, doi:10.1038/s41467-017-00524-5 (2017).

47 Acemel, R. D., Maeso, I. & Gomez-Skarmeta, J. L. Topologically associated domains: a successful scaffold for the evolution of gene regulation in animals. Wiley Interdiscip Rev Dev Biol 6, doi:10.1002/wdev.265 (2017).

## Methods’ References

48 Moreno-Mateos, M. A. et al. CRISPRscan: designing highly efficient sgRNAs for CRISPR-Cas9 targeting in vivo. Nat Methods 12, 982–988, doi:10.1038/nmeth.3543 (2015).

49 Carmona-Aldana, F. et al. CTCF knockout reveals an essential role for this protein during the zebrafish development. Mech Dev 154, 51–59, doi:10.1016/j.mod.2018.04.006 (2018).

50 Freitas, R., Gomez-Marin, C., Wilson, J. M., Casares, F. & Gomez-Skarmeta, J. L. Hoxd13 contribution to the evolution of vertebrate appendages. Dev Cell 23, 1219–1229, doi:10.1016/j.devcel.2012.10.015 (2012).

51 Tena, J. J. et al. Odd-skipped genes encode repressors that control kidney development. Dev Biol 301, 518–531, doi:10.1016/j.ydbio.2006.08.063 (2007).

52 Dobin, A. et al. STAR: ultrafast universal RNA-seq aligner. Bioinformatics 29, 15–21, doi:10.1093/bioinformatics/bts635 (2013).

53 Anders, S., Pyl, P. T. & Huber, W. HTSeq--a Python framework to work with high-throughput sequencing data. Bioinformatics 31, 166–169, doi:10.1093/bioinformatics/btu638 (2015).

54 Love, M. I., Huber, W. & Anders, S. Moderated estimation of fold change and dispersion for RNA-seq data with DESeq2. Genome Biol 15, 550, doi:10.1186/s13059-014-0550-8 (2014).

55 Huang da, W., Sherman, B. T. & Lempicki, R. A. Systematic and integrative analysis of large gene lists using DAVID bioinformatics resources. Nat Protoc 4, 44–57, doi:10.1038/nprot.2008.211 (2009).

56 Buenrostro, J. D., Giresi, P. G., Zaba, L. C., Chang, H. Y. & Greenleaf, W. J. Transposition of native chromatin for fast and sensitive epigenomic profiling of open chromatin, DNA-binding proteins and nucleosome position. Nat Methods 10, 1213–1218, doi:10.1038/nmeth.2688 (2013).

57 Fernandez-Minan, A., Bessa, J., Tena, J. J. & Gomez-Skarmeta, J. L. Assay for transposase-accessible chromatin and circularized chromosome conformation capture, two methods to explore the regulatory landscapes of genes in zebrafish. Methods Cell Biol 135, 413–430, doi:10.1016/bs.mcb.2016.02.008 (2016).

58 Santos-Pereira, J. M., Gallardo-Fuentes, L., Neto, A., Acemel, R. D. & Tena, J. J. Pioneer and repressive functions of p63 during zebrafish embryonic ectoderm specification. Nat Commun 10, 3049, doi:10.1038/s41467-019-11121-z (2019).

59 Schmidl, C., Rendeiro, A. F., Sheffield, N. C. & Bock, C. ChIPmentation: fast, robust, low-input ChIP-seq for histones and transcription factors. Nat Methods 12, 963–965, doi:10.1038/nmeth.3542 (2015).

60 Langmead, B. & Salzberg, S. L. Fast gapped-read alignment with Bowtie 2. Nat Methods 9, 357–359, doi:10.1038/nmeth.1923 (2012).

61 Li, H. et al. The Sequence Alignment/Map format and SAMtools. Bioinformatics 25, 2078–2079, doi:10.1093/bioinformatics/btp352 (2009).

62 Quinlan, A. R. & Hall, I. M. BEDTools: a flexible suite of utilities for comparing genomic features. Bioinformatics 26, 841–842, doi:10.1093/bioinformatics/btq033 (2010).

63 Haeussler, M. et al. The UCSC Genome Browser database: 2019 update. Nucleic Acids Res 47, D853–D858, doi:10.1093/nar/gky1095 (2019).

64 Zhang, Y. et al. Model-based analysis of ChIP-Seq (MACS). Genome Biol 9, R137, doi:10.1186/gb-2008-9-9-r137 (2008).

65 Ramirez, F. et al. deepTools2: a next generation web server for deep-sequencing data analysis. Nucleic Acids Res 44, W160–165, doi:10.1093/nar/gkw257 (2016).

66 Heinz, S. et al. Simple combinations of lineage-determining transcription factors prime cis-regulatory elements required for macrophage and B cell identities. Mol Cell 38, 576–589, doi:10.1016/j.molcel.2010.05.004 (2010).

67 Hiller, M. et al. Computational methods to detect conserved non-genic elements in phylogenetically isolated genomes: application to zebrafish. Nucleic Acids Res 41, e151, doi:10.1093/nar/gkt557 (2013).

68 Li, H. & Durbin, R. Fast and accurate short read alignment with Burrows-Wheeler transform. Bioinformatics 25, 1754–1760, doi:10.1093/bioinformatics/btp324 (2009).

69 Durand, N. C. et al. Juicer Provides a One-Click System for Analyzing Loop-Resolution Hi-C Experiments. Cell Syst 3, 95–98, doi:10.1016/j.cels.2016.07.002 (2016).

70 Knight, P. A. & Ruiz, D. A fast algorithm for matrix balancing. IMA Journal of Numerical Analysis 33, 1029–1047, doi:10.1093/imanum/drs019 (2012).

71 Kruse, K., Hug, C. B. & Vaquerizas, J. M. FAN-C: A Feature-rich Framework for the Analysis and Visualisation of C data. bioRxiv, 2020.2002.2003.932517, doi:10.1101/2020.02.03.932517 (2020).

72 Lieberman-Aiden, E. et al. Comprehensive mapping of long-range interactions reveals folding principles of the human genome. Science 326, 289–293, doi:10.1126/science.1181369 (2009).

73 Schwartzman, O. et al. UMI-4C for quantitative and targeted chromosomal contact profiling. Nat Methods 13, 685–691, doi:10.1038/nmeth.3922 (2016).

